# Inference of human pigmentation from ancient DNA by genotype likelihood

**DOI:** 10.1101/2025.01.29.635495

**Authors:** Silvia Perretti, Maria Teresa Vizzari, Patrícia Santos, Enrico Tassani, Andrea Benazzo, Silvia Ghirotto, Guido Barbujani

**Author notes:** **Author Contributions:** SP analyzed the data and wrote the manuscript. MTV, PS, ET analyzed the data. AB contributed new analytic tools. SG and GB designed research and wrote the manuscript. **Competing Interest Statement:** The authors declare no competing interest.

## Abstract

Light eyes, hair and skins probably evolved several times as *Homo sapiens* dispersed from Africa. In areas with lower UV radiation, light pigmentation alleles increased in frequency because of their adaptive advantage and of other contingent factors such as migration and drift. However, the tempo and mode of their spread is not known. Phenotypic inference from ancient DNA is complicated, both because these traits are polygenic, and because of low sequence depth. We evaluated the effects of the latter by randomly removing reads in two high-coverage ancient samples, the Paleolithic Ust’-Ishim from Russia and the Mesolithic SF12 from Sweden. We could thus compare three approaches to pigmentation inference, concluding that, for suboptimal levels of coverage (<8x), a probabilistic method estimating genotype likelihoods leads to the most robust predictions. We then applied that protocol to 348 ancient genomes from Eurasia, describing how skin, eye and hair color evolved over the past 45,000 years. The shift towards lighter pigmentations turned out to be all but linear in time and place, and slower than expected, with half of the individuals showing dark or intermediate skin colors well into the Copper and Iron ages. We also observed a peak of light eye pigmentation in Mesolithic times, and an accelerated change during the spread of Neolithic farmers over Western Eurasia, although localized processes of gene flow and admixture, or lack thereof, also played a significant role.

## Introduction

Skin is not preserved in fossils, but there is little doubt that the early hominins’ epidermis was covered by hair and lightly pigmented (1). Exposure to sunlight induces degradation of folate, a molecule essential in DNA synthesis and cell proliferation (2); as the hair protection was lost in the course of evolution, there is evidence for a selective sweep favoring alleles that would make skins darker (3). Conversely, light skin colors promote photochemical-controlled synthesis of vitamin D (2). When *Homo* spread Northwards from Africa into Eurasia, the selection regime thus changed, and lighter phenotypes emerged. In both phases UV radiation seems the most likely selective agent, since other environmental factors such as temperature, rainfall or humidity show lower levels of correlation with skin color (4). On top of these major trends, adaptation to local conditions, genetic drift and migration contributed to past and present patterns of human pigmentation.

Skin, eye and hair colors are complex phenotypes with polygenic inheritance. All depend on the different amounts, type, and distribution of two pigments, eumelanin (brown-red) and pheomelanin (brown-yellow), produced by human melanocytes (5). Blue and green irises are not due to additional eye pigments, but to the scattering of the light caused by variable cellular density of the corneal stroma (5).

The melanin biosynthetic pathway involves several enzymatic steps, and at least 26 relevant genes have been identified in association studies (6). As is the case for other complex traits, AI-based algorithms have been designed to predict skin, eye and hair color from DNA information. One of the most widely used test systems, HIrisPlex-S (7–9), infers from 41 SNPs the individual probabilities for three eye, four hair, and five skin color categories. When based on good-quality genomic data, HIrisPlex-S has a low margin of error. However, problems may arise when, regardless of the inferential method used, phenotypes are to be inferred from ancient genomes, usually typed at low coverage.

HIrisPlex-S assumes that the allelic and genotypic states of the loci of interest are known, which for many ancient samples is unwarranted. Indeed, directly calling genotypic variants from such data is challenging, because of DNA fragmentation, exogenous contamination, and degradation. More often than not, all these factors result in low sequencing depth, which in turn affects the possibility to reliably identify genotypes, and hence phenotypes too. Thus, the robustness of phenotypic prediction methods when applied to ancient samples needs to be validated considering their particular characteristics, as also remarked by Manuel Ferrando-Bernal *et al*., 2025 (10).

A critical factor, in these cases, is the possibility to take into account genotype uncertainty. In the first part of this paper we compared three approaches to pigmentation inference, concluding that, for suboptimal levels of coverage (<8x), a probabilistic method estimating genotype likelihoods leads to the most robust predictions. Then, in the second part of this paper, we applied that protocol to a broad dataset of 348 ancient genomes from Eurasia, thus describing how skin, eye and hair color evolved over the past 45,000 years.

## Results

### Testing the robustness of phenotypic inference on ancient data

The primary objective of this study was to evaluate the effect of different levels of genome coverage on phenotypic inference in the context of low-coverage ancient DNA studies. HIrisPlexS was developed in forensic science, but that has been recently used even in the context of ancient DNA studies. The HIrisPlex-S protocol has been applied on various ancient specimens, including, for example, medieval samples (11), remains dated to around 1485 attributed to King Richard III (12), the “Cheddar Man” (13), the “La Braña” Mesolithic sample (14), and even a 5700-year-old chewed birch pitch from which human DNA was extracted (15). So far, however, no one tested whether the protocol is effective on samples sequenced at low coverage, or whether it leads to robust inference when different calling algorithms are used. As detailed in Figure 1, we evaluated the HIrisPlex-S inferential robustness with genotypes generated using three different calling procedures: a *direct* approach, in which genotypes are directly called using *GATK UnifiedGenotyper* v3.5 (16); an *imputation* protocol, commonly used in the ancient DNA context, performed using *GLIMPSE* v1.1.1 (17); and a *probabilistic* approach, where genotypes likelihoods (18) were calculated and 1,000 different genotypes were sampled, weighting their effect on the phenotypic inference according to their likelihood. We obtained phenotypic prediction by analyzing the 41 HIrisPlexS positions: direct and imputed genotypes were directly entered into the HIrisPlex-S website, whereas probabilistic predictions incorporated likelihood-based genotype sampling. We then postprocessed the resulting prediction files in *R* environment v4.3.3 (19) to extract, interpret, and compare the phenotypic outcomes across all approaches (see Methods and SI for details).

**Figure 1.**
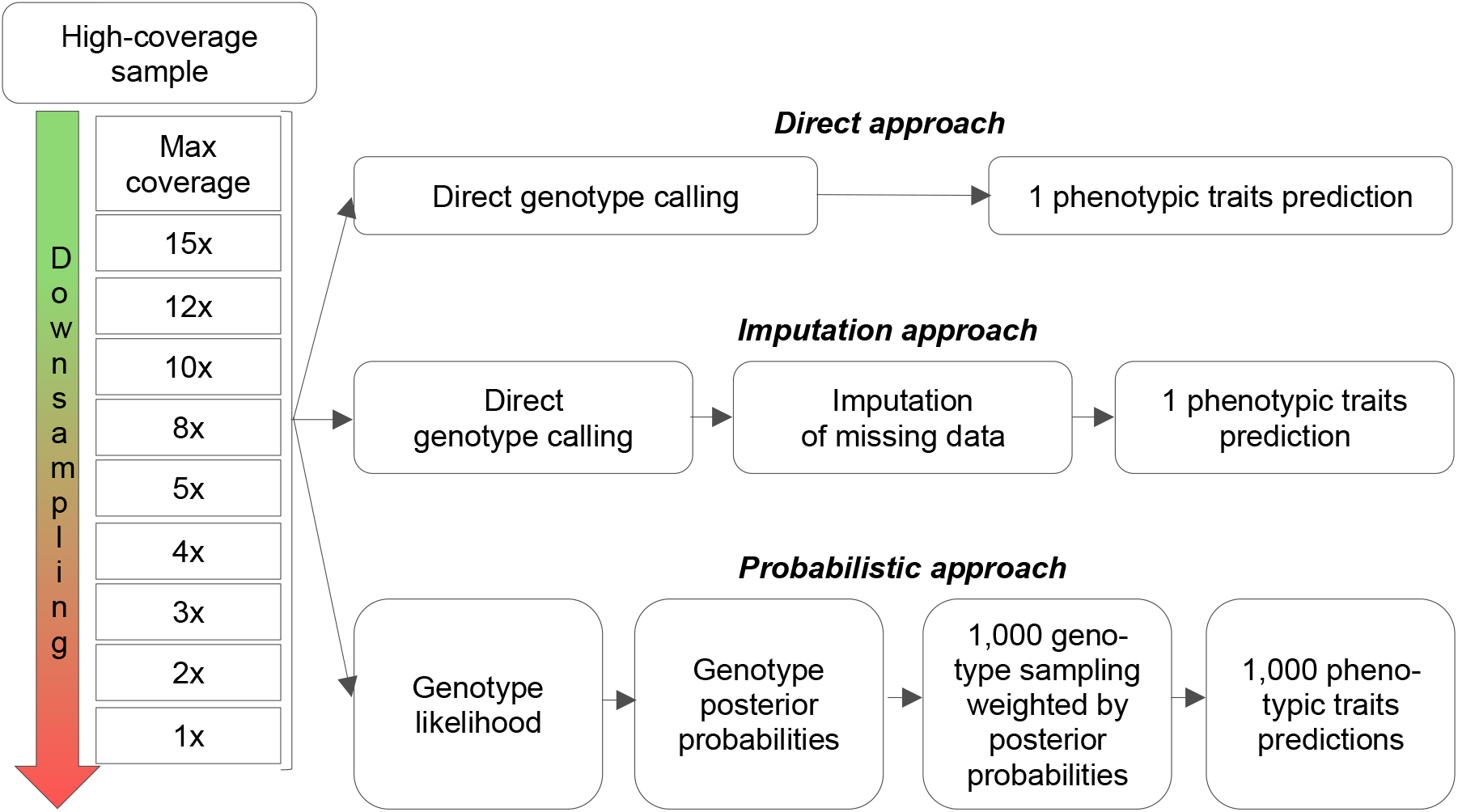
Diagram illustrating the conceptual framework for phenotypic prediction across varying sequencing coverages. The original high-coverage sample (top) undergoes progressive downsampling, decreasing sequencing coverage (left). The downsampled data are processed through three distinct workflows: direct approach (top branch), imputation approach (middle branch), and probabilistic approach (bottom branch). Each branch represents a step-by-step workflow that returns predicted phenotypic traits for downstream comparative analyses.

To measure the loss of information when coverage is low, we iteratively downsampled the Ust’-Ishim (45,045 calBP from western Siberia, (20)), and SF12 (9,033±8,757 calBP from Sweden, (21)) samples’ reads. We selected these samples because they represent crucial historical periods and geographical locations (Paleolithic Russia and Mesolithic Sweden), and were, at the time of the analysis, the two best-covered samples at the 41 HIrisPlex-S positions.

We first obtained the phenotypic prediction for each sample using the high-coverage genome (28x for the Ust’-Ishim sample and 44x for the SF12 sample). Subsequently, we identified the minimum coverage across the 41 HIrisPlex-S positions, that is 17x for Ust’-Ishim sample and 33x for SF12, and performed progressive downsampling starting from these coverage levels. We thus compared the phenotypic predictions obtained from the high-coverage genomes with the ones obtained considering 10 or 11 different coverage levels: 17x, 15x, 12x, 10x, 8x, 5x, 4x, 3x, 2x, 1x for the Ust’-Ishim sample and 33x, 20x, 15x, 12x, 10x, 8x, 5x, 4x, 3x, 2x, 1x for the SF12 sample. For each coverage level, we tested 10 independent downsampled datasets. Figure 2 shows the estimated phenotypes and their frequency for each coverage level for the two test samples, under the three different calling algorithms.

**Figure 2.**
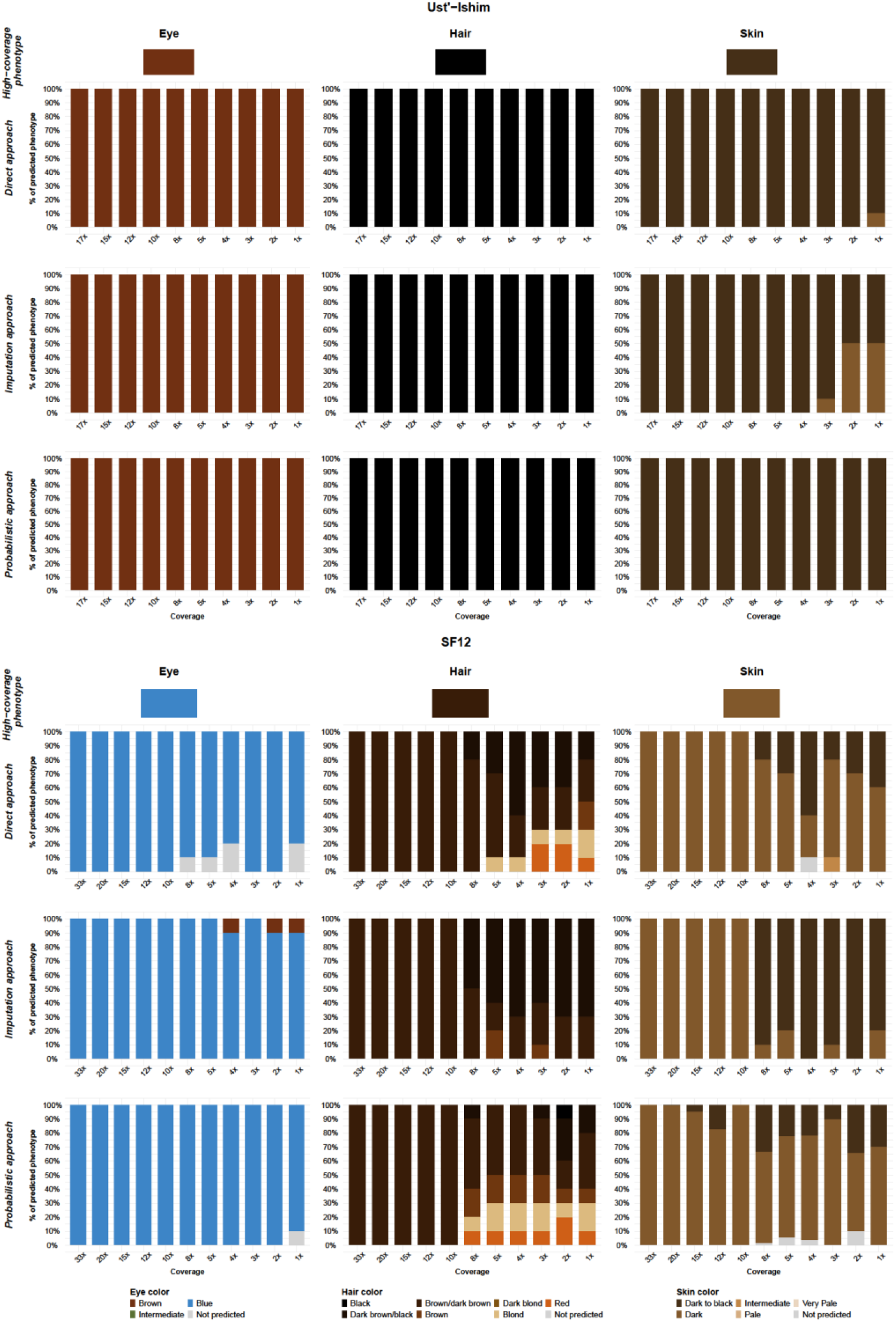
Comparative phenotypic predictions for Ust’-Ishim (Panel A) and SF12 (Panel B) across the 3 different approaches and sequencing coverages. Columns correspond to the three different phenotypic traits predicted: eye color (left), hair color (center), and skin color (right). Each panel consists of four rows: the first row presents the phenotypes inferred from high-coverage data (17x for Ust’-Ishim and 33x for SF12 respectively) following the direct approach. The second, third, and fourth rows show the results obtained using the direct, imputation, and probabilistic approaches, respectively. The x-axis represents the different coverage levels, while the y-axis shows the percentage of times each phenotype was predicted over 10 iterations.

#### Ust’-Ishim sample

what we might call the *true* phenotypes were evaluated through the *direct* approach with a mean coverage level of 28x. The HIrisPlex-S system estimated brown eyes, black hair and dark to black skin. As it is shown in Figure 2A, for this Paleolithic sample the three estimation methods returned robust estimates for eyes and hair color, regardless of the coverage level, correctly predicted 100% of times. Conversely, the skin prediction yielded different results. The *true* phenotype was recognized all times only through the *probabilistic* approach; with the *direct* approach we had 10% of error in the estimate at the lowest coverage levels, whereas the *imputation* protocol started providing erroneous estimates at 3x (10% of error), reaching the 50% of wrong predictions at 2 and 1x.

#### SF12 sample

the *true* phenotypes were evaluated through the *direct* approach with a mean coverage level of 44x, namely blue eyes, brown hair and dark skin (Fig. 2B). With the *direct* approach the system was unable to predict the eye color for some downsampled datasets starting at 8x, whereas with the *probabilistic* approach the eye color was unpredicted only in the 10% of the 1x datasets. In all the other downsampled datasets the blue eyes phenotype was correctly identified. With the *imputation* approach the system failed to identify the *true* phenotype at 4, 2 and 1x, where in 10% of cases brown eyes were inferred. For this sample the hair color was correctly predicted with greater difficulty for all the three approaches tested. We indeed observed wrong estimation starting at 8x of coverage, with a misassignment proportion reaching 80% for some coverage levels. Among the three approaches tested, the *probabilistic* method was the one showing the lowest misidentification rate, whereas the highest error rate was observed for the imputation *approach* (68% of misidentification rate on average below 8x). The skin color was correctly predicted 100% of times with the three approaches when the coverage level was equal to or higher than 10x. Starting at 8x the system showed some issues in correctly predicting the *true* phenotype, especially when the genotypes were defined through the *direct* approach or *imputation* method. In particular, datasets generated through the *imputation* method were misassigned almost 100% of times for coverage levels below 8x. This percentage is lower when the *direct* approach is used, about 30% on average. The *probabilistic* approach performed slightly better, below a 8x of coverage the percentage of misassigned dataset is 23% on average.

This analysis highlights how easily the HIrisPlex-S phenotypic estimation procedure fails, if genotypes are called from data with a coverage equal to or lower than 8x, a situation commonly encountered when DNA comes from ancient specimens. Above this coverage threshold, there are no significant differences between the *direct, probabilistic* and *imputation* methods; the *true* phenotype is recovered 100% of times. At a genomic coverage level of 8x or lower, conversely, the three methods perform differently. Among them, the *probabilistic* method is the one that most frequently returns the same phenotype estimated at high coverage. The worst performances are provided by the *imputation* method, suggesting that the phenotypes inferred obtained through imputation of all missing HIrisPlex-S positions should be considered with caution.

### Phenotypic inference of Eurasian samples from Paleolithic to Iron Age

We collected a large dataset of 348 published ancient DNAs from individuals typed at different coverage levels (all above 1x; Fig. 3 and S1_Appendix), and spanning from 45,000 to 1,700 years ago. Samples were labelled based on archaeological evidence and not only on chronology. We estimated eye, skin, and hair color of each sample, applying the inferential approach that provided more robust results for that specific coverage level in the previous analyses (see Methods and Supplementary Appendix files from S2 to S7). Figure 4 and Supplementary figures S13-S24 show the pigmentation traits inferred in samples belonging to different time periods. We only reported phenotypes that the calling methods could confidently predict; for the *probabilistic* method we reported the status of a phenotype only if it had been predicted in at least 90% of the 1,000 replications generated by the genotype likelihoods.

**Figure 3.**
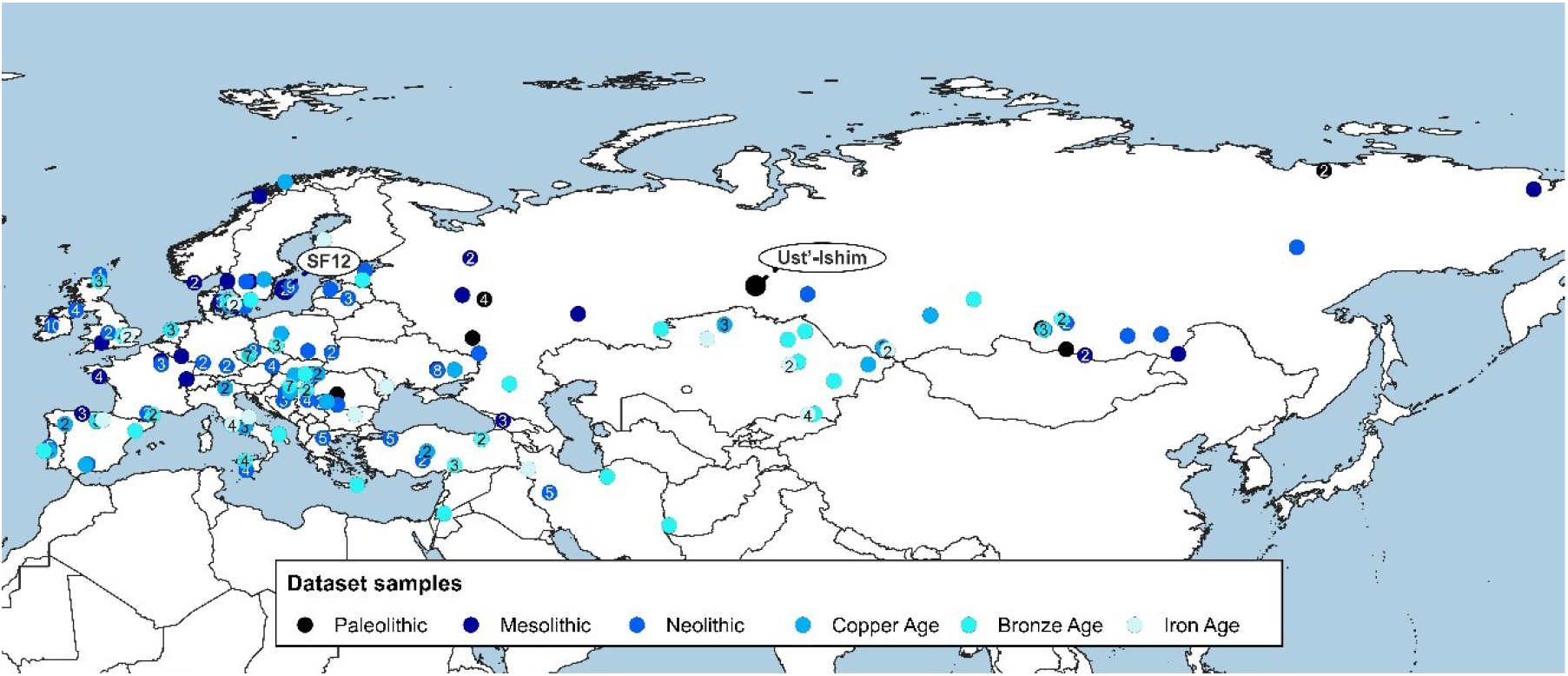
Geographical distribution across Eurasia of selected ancient samples. The colored dots represent different time transect: black-Paleolithic, light blue-Iron Age, and intermediate shades-transitional periods (from Mesolithic to Bronze Age). The number of samples at each site is indicated inside each dot. The two test samples, the Paleolithic Ust’-Ishim and the Mesolithic SF12, are highlighted with labels.

**Figure 4.**
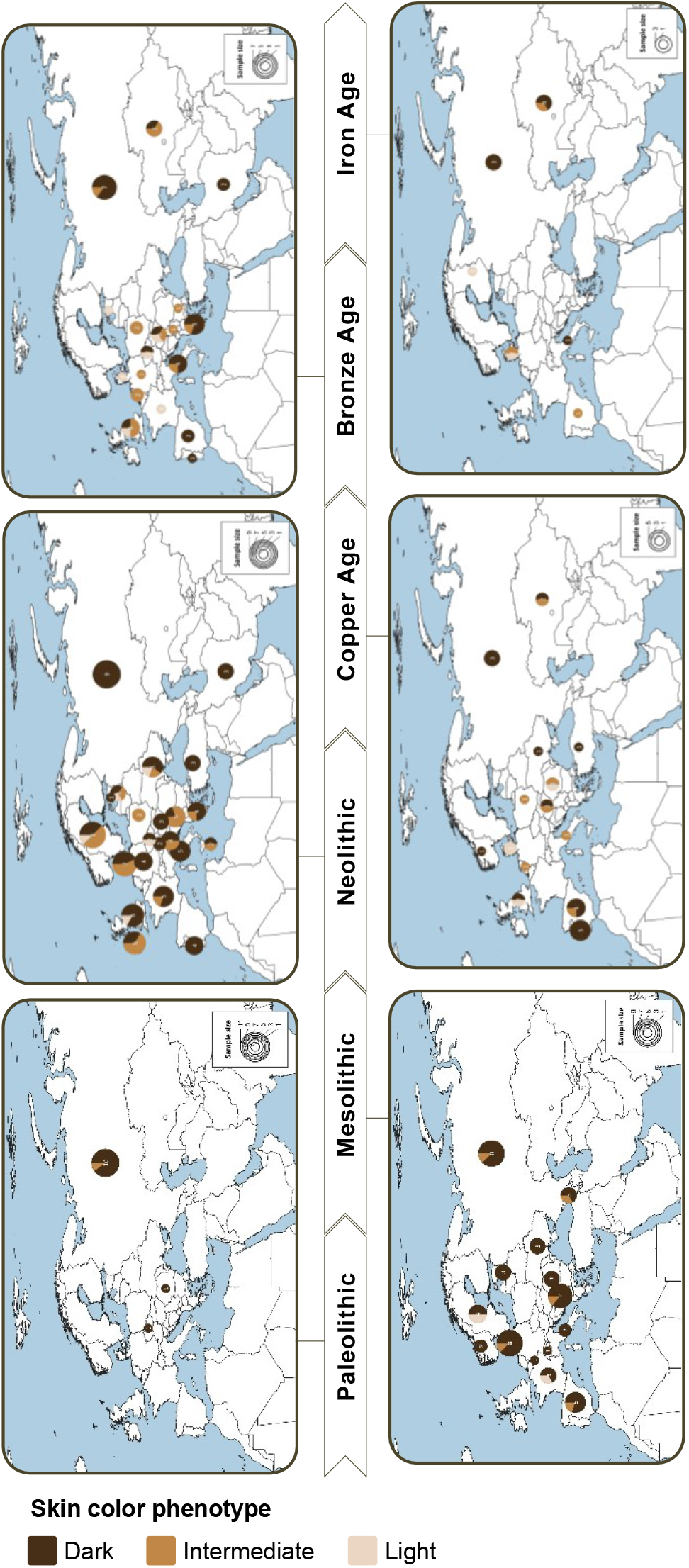
Temporal and geographical distribution of skin pigmentation estimates in Eurasia from Paleolithic to Iron Age. The maps illustrate the spatial and temporal distribution of the inferred skin pigmentation phenotypes. Dimension of each pie chart indicates the sample size. Skin color results are grouped into three categories: Dark (Dark to Black and Dark), Intermediate, and Light (Pale and Very Pale).

#### Paleolithic period

(from approximately 45,000 to 13,000 years ago; 12 samples; 11 typed for eye color, hereafter E, 10 for hair color, hereafter H; 12 for skin color, hereafter S; one of them is the Ust’-Ishim test sample). Dark phenotypes are inferred for all traits for almost all the samples analyzed. The only exception is a Russian sample, Kostenki 14, dated to between 38,700 and 36,200 years ago, which exhibits an intermediate skin color (22).

#### Mesolithic period

(from approximately 14,000 to 4,000 years ago; 66 samples; 35 E, 63 H, 53 S; one of them is the SF12 test sample). Light eye colors are inferred for 11 samples; they come from Northern Europe, France and Serbia. By contrast, all 24 samples from the easternmost regions only display the dark phenotype. In Serbia both phenotypes coexist, one with blue eyes and four with brown eyes. 61 samples show dark hair phenotypes, with the exception of 1 Swedish and 1 Serbian sample, both showing blonde features. Skin color displays a broader range of phenotypes: predominantly dark (43 samples), with regions in Europe also showing intermediate phenotypes (seven samples from Denmark, France, Georgia, Russia, Serbia, and Spain) and the earliest light phenotypes observed in this study (three samples from France and Sweden). In this time transect we observe for the first time an individual with inferred blue eyes, blonde hair, and light skin, NEO27, a hunter-gatherer from Sweden who lived approximately 12,000 years ago (23).

#### Neolithic period

(from approximately 10,000 to 4,000 years ago; 132 samples, 93 E, 120 H, 93 S). We still observe the majority of individuals showing the dark eye phenotype (81 samples), including France, in which we previously found only light phenotype. Both dark and light eye phenotypes are observed in Northern and Central-Eastern Europe, with the light phenotype inferred in 12 samples from Austria, Denmark, Greece, Ireland, Latvia, Serbia, and Sweden. Hair color is predicted as dark in almost all samples, with one exception from Austria who has an intermediate phenotype and five from Denmark, Greece, Ireland, and Serbia with light phenotype. Additionally, we observed for the first time in our dataset one sample with red hair, from Turkey. The skin phenotype is more variable, with regions in Europe (Portugal, Italy, Austria, Germany, Hungary, Estonia, and Russia) and Western Asia (Iran and Turkey) exhibiting exclusively a dark phenotype, whereas other regions show either both dark and intermediate phenotypes (25 samples exhibit the latter, from Croatia, Denmark, France, Greece, Ireland, Latvia, Malta, Poland, Serbia, Sweden, and Ukraine), or even light skin phenotypes (in five samples from the Czech Republic, Great Britain, Latvia, Sweden, and Ukraine).

#### Copper Age

(from approximately 6,000 to 3,500 years ago; 42 samples, 31 E, 33 H, 28 S). Even during the Copper Age dark phenotypes are prevalent. Most samples, 26, showed dark eyes, with the light phenotype present in five samples from Denmark, Hungary, Italy, and Romania.

Hair phenotypes remain mostly dark, with only one sample showing intermediate hair color (Denmark) and one samples exhibiting light hair color (Romania). Skin color is still predominantly dark (17 samples) in Eastern Europe, and the Iberian Peninsula, but intermediate skin tones are observed in Spain, Kazakhstan, and Central Europe (seven samples from Hungary, Italy, the Netherlands, Poland, and Romania), and light skins in Denmark, Great Britain, and Romania (four samples).

#### Bronze Age

(from approximately 7,000 to 3,000 years ago; 71 samples, 55 E, 64 H, 43 S). In this time period we observed an increasing proportion of light eye phenotype. While 39 samples throughout Europe and Asia are still exhibiting dark eyes, 16 samples display a light phenotype. These light phenotypes are still mainly found in Europe, but are also emerging in other regions such as Russia and Jordan, and as far East as Kazakhstan. Dark hair phenotypes remain predominant in most of Europe and Asia (49 samples), with intermediate phenotypes present in two samples from Denmark and Hungary. However, there is a greater proportion of light phenotypes (12 samples), specifically in Northern and Central-Eastern Europe, and they appear in Italy, Russia, Jordan, and Kazakhstan. One sample from Greece exhibits red hair. Western Europe, Southern Europe, Russia, and Southern Asia still exhibit a higher frequency of dark skin phenotypes (22 samples), but we also observed an increase in intermediate phenotypes in Central Europe and Central-Eastern Europe, as well as their first appearance in Russia (15 samples in total). The light phenotype emerged in six samples from the Czech Republic, Denmark, Estonia, France, Great Britain, and Hungary. During this period, we observed an increase in the co-occurrence of estimated blue eyes, blonde hair, and light skin, with four samples exhibiting this combination of phenotypes: I7198 from the Czech Republic (14), EKA1 from Estonia (24), I2445 from England (14), and SZ1 from Hungary (25).

#### Iron Age

(from approximately 3,000 to 1,700 years ago; 25 samples, 15 E, 19 H, 11 S). In this phase, the dark eye phenotype (10 samples) is present in Great Britain, Spain, and Russia, while the light eye phenotype (3 samples) is found in Denmark and Finland. Italy, and Kazakhstan exhibit both phenotypes. Hair remains predominantly dark throughout Europe and Asia (14 samples), with one intermediate phenotype observed in Denmark and four light phenotypes in Denmark, Finland, Italy, and Kazakhstan. Skin color analysis shows the dark phenotype (six samples) in Russia, Kazakhstan, and Italy. The intermediate phenotype (three samples) in Denmark, Kazakhstan, and reappears in Spain. The light phenotype (two samples) is still present in Northern Europe. A combination of blue eyes, blonde hair, and pale skin is observed in two samples: VK521 from Denmark (26) and DA236 from Finland (27).

## Discussion

Inference of human pigmentation traits from ancient DNA evidence has garnered significant attention in recent years. This inferential procedure aims to reconstruct past phenotypic characteristics by examining genetic polymorphisms associated with eye, hair and skin color. So far, however, all the available approaches rest on the assumption that the sample’s allelic state at polymorphic positions is exactly known, which is not always warranted. Indeed, ancient genomic data are often produced at low coverage, meaning that often few reads (if any) map a specific genomic region; as a consequence, the direct calling of genotypes on these samples may introduce a substantial bias in downstream analyses. The HIrisPlex-S system proved to be very efficient in correctly reconstructing human pigmentation traits (7–9). However, our downsampling experiments show that it can be safely exploited through the currently available pipeline (*direct* approach) only when the minimum coverage level at the 41 inferential sites is at least 8x, which is uncommon in ancient samples.

To address this issue, in this work we propose a framework integrating a *probabilistic* approach in phenotype inference, useful when a direct genotype calling would not be accurate. We tested this framework by estimating phenotypes considering for each sample 1,000 combinations of genotypes at the 41 HIrisPlex-S positions, reflecting their likelihoods (18). Through this *probabilistic* approach we obtain both a phenotype inference and a measure of its reliability. The coverage downsampling procedure we applied on the Ust’-Ishim and SF12 samples proved the probabilistic approach is robust in inferring the *true* pigmentation trait even at low (below 8x) or very low (1-2x) coverage levels. With low-coverage data it is quite common to have positions that are not covered by any reads. In such cases, one approach, widely used in paleogenomics, is to impute the genotypic state using algorithms that leverage information from adjacent loci and a reference database to fill in the missing position. However, we showed that imputing all the 41 HIrisPlex-S positions may bias the estimation of phenotypes, particularly at very low-coverage, and therefore the *imputation* approach should be taken with caution when aiming at phenotypic inference. As is the case when continuous traits are described as discrete, not all phenotypes are inferred with the same level of confidence by HIrisPlex-s (7–9). This is particularly true for the skin phenotypes categories, the limits of which are admittedly somewhat arbitrary.

By a *probabilistic* approach, we showed that eye, hair and skin color changed substantially through time in Eurasia. It was reasonable to imagine that the first hunting-gathering settlers, who came from warmer climates, had mostly dark pigmentation (2). What was less expected was the long persistence of their phenotypes. We found the first instance of light skin color in the Swedish Mesolithic, but it comes from only one sample in >50. Things changed afterwards, but very slowly, so that only in the Iron Age did the frequency of light skins equal that of dark skins; during much of prehistory, most Europeans were dark-skinned. A similar trend, with dark pigmentation long coexisting with an increasing, yet relatively small proportion of lighter traits, is observed for hair and eye color, although there was a temporary peak of light eye frequency in the Mesolithic period, when we inferred light pigmentation for 11 out of 35 samples.

There is little doubt that, on top of selection, gene flow was the main factor causing shifts of pigmentation traits. Antonio *et al*. (2024) observed a decrease of between-population genetic variances as we move in time from the Paleolithic period to the present (28). In this study, we described a parallel increase of within-population variance for pigmentation traits. Both results are the expected consequences of an evolutionary scenario in which the effects of gene flow exceed those of genetic drift.

Among the episodes of gene flow documented in Western Eurasian prehistory, the spread of early Neolithic farmers from Anatolia is known to have profoundly changed the genetic makeup of populations (29), to the point that some authors speak of ‘population turnover’ e.g. in the British Isles (13) and in Denmark (23). What we observed for pigmentation traits appears to be, in large part, a consequence of that massive migration. Actually, the transition to food production also led to an increase in infectious disease and to a poorer diet (see e.g. Pearson *et al*., 2023) (30), but once in the new territories, immigrating farmers had two evolutionary advantages over their hunting-gathering counterparts. By farming and animal herding they increased the amount of available food, and had a skin phenotype fit for the lower levels of UV radiation. Both factors gave them (and only them) the potential for demographic growth (31, 32), ultimately leading to profound changes in the Europeans’ genomes.

However, these and other ancient DNA data (33) show that the process was all but linear, and took longer than the Neolithic to be completed. The first instance of light phenotypes we identified dates back to Mesolithic (skin, eye) or Neolithic (hair) times, but whereas light skins steadily increased in frequency across time, possibly due to their adaptive value, hair and eye colors showed fluctuations, the main of which is the localized Mesolithic increase of light eye frequency. As a consequence, we do not think that the changes described in this paper can be regarded as the effects of a wave of migration proceeding at a regular pace. Rather, what we think we are observing is a process in which, above and beyond the major Neolithic demic diffusion over much of Western Eurasia, localized processes of migration and admixture, or lack thereof, played a significant role.

Identifying the specific genes responsible for the observed trends is complicated due to the low mean coverage, which often leaves the allelic status at several loci undefined. However, comparing the Paleolithic sample Ust-Ishim (with dark phenotypes for all 3 traits) to the Hungarian Bronze age SZ1 (with light phenotypes for all 3 traits) reveals allelic differences at several loci known to have changed in the course of time *(SLC45A2, LOC105374875, HERC2, PIGU, TYRP1, ANKRD11, BNC2, HERC2, OCA2, DEF8*), as well as two novel loci, *TYR* (rs1042602) and *SLC24A5* (rs1426654), both of which carry the AA genotypes in the SZ1 sample. Strong support from previous studies links the A allele of rs1042602 to the absence of freckles (34). Furthermore, the rs1042602 variant, located in the *TYR* gene encoding tyrosinase involved in melanin biosynthesis, is associated with albinism, particularly in individuals who are homozygous for the A allele, in combination with another genetic variant within the *TYR* gene, rs4547091, when homozygous for the C allele (35). The nonsynonymous mutation (G -> A) at rs1426654 in the *SLC24A5* gene is strongly associated with reduced melanin production and lighter skin pigmentation in humans. The ancestral allele G is fixed in Africans and East Asians while the derived allele A is nearly fixed in European populations (98.7-100%) (36, 37) or present at high in frequencies in Near East and South Asian populations. In addition, rs1426654 has been linked to skin pigmentation variation in admixed populations with recent European ancestry and within South Asians (38). Its rarity in East Asia and most sub-Saharan African populations supports the hypothesis that it originated in Europe or the Near East within the past 10,000 to 35,000 years (38). As noted by Lin *et al*. in their 2018 study, the *SLC24A5* locus exemplifies a rare instance of strong, recent adaptation in human history and serves as a case of adaptive gene flow at a pigmentation-related locus (38).

HIrisPlex-S has a low margin of error particularly for European populations. While the system provides robust inferences, its accuracy can be affected in recently admixed populations, especially for intermediate phenotypes (39, 40) and skin color. This can be attributed to factors such as phenotype misclassification resulting from subjective skin tone perception, as observed by Carratto *et al*., 2019 in Southeastern Brazilians (41). With the development of inferential procedures for phenotype estimation, such as the one proposed in this work, together with the increasing production of high-quality ancient genomic data, we will gain unprecedented insights into the evolution of pigmentation-related phenotypes in our species. This will help reconstruct a comprehensive and detailed picture of crucial phases of our evolutionary history while enhancing our understanding of the evolutionary and demographic forces that drove and shaped modern human phenotypic variation.

## Methods

### Testing the robustness of phenotypic inference on ancient data

To evaluate the robustness of the three different approaches we selected two ancient highcoverage samples, Ust’-Ishim and SF12. For the *direct* and *probabilistic* approaches we generated a *pileup* file from the alignment data for the 41 HIrisPlex-S positions using *SAMtools* v1.11 *mpileup* command (42). Then, we conducted a pointwise progressive downsampling from the *mpileup* output file within the *R* environment v4.3.3 (19), using a non-replacement sampling to ensure random selection of unique elements to prevent read and nucleotide duplication (see SI for details). For the *imputation* approach the downsampling was performed using the *SAMtools* v1.11 *view* command with the *-s* option (42), where the 41 informative positions have been masked and then imputed (see SI for details).

For the *probabilistic* approach, we computed, for each of the 41 informative positions, the genotype likelihoods for each of the ten possible genotypes within the *R* environment v4.3.3 (19) applying the formula of the first version of *GATK* (dragon) (43). This approach ensures that multiple genotypes may be evaluated for the same position due to their genotype likelihood values. Since the HIrisPlex-S system accepts only a single allele derived from a genotype, we performed 1,000 samplings from the 10 possible genotypes according to their genotype likelihoods and posterior probabilities. Each sampling event resulted in a unique combination of informative alleles. The final result consists of a table containing 1,000 rows, each representing a set of 41 allelic combinations (see SI for details).

To infer the phenotypic traits, we uploaded the required *CSV* file containing the allele count for the 41 positions, obtained with the three protocols, into the HIrisPlex-S website (https://hirisplex.erasmusmc.nl). The results obtained from the HIrisPlex-S system were analyzed within the *R* environment v4.3.3 (19) following the guidelines outlined in the HIrisPlex-S User Manual (2018) (7–9).

### Inference of pigmentation traits on ancient genomic data

We analyzed a dataset of 348 published ancient human whole-genomes (Fig. 3 and S1_Appendix), including the two test samples used for the validation step, with a minimum mean coverage of 1x. The samples encompass a temporal range from approximately 45,000 to 1,700 years ago, and distributed from Western Europe to Asia, representing a total of 34 countries. We defined six different groups based on the chronological period, archeological context, cultural affiliation and genetic affinities of the samples, as reported in the literature. All the samples were processed using an *in-house* pipeline (see SI for details).

Within our dataset, 13 samples have a coverage level above 8x across all the 41 HIrisPlexS positions, and so we applied the *direct* approach to extract the genotypes for the 41 positions; 335 samples for which the coverage was equal to or below 8x we applied the *probabilistic* approach and we treated uncovered HIrisPlex-S positions as missing data.

All data and code underlying the inferential procedure presented in this manuscript are fully documented and available on GitHub at https://github.com/Ghirotto-Lab-at-University-of-Ferrara/Phenotypic_inference.

## Acknowledgments

GB and SP were financially supported by PRIN 2020 (grant 2020HJXCK9) from the Italian Ministry of Education, University and Research (MIUR). MTV, AB, PAS and SG were financially supported by PRIN 2020 (grant 2020TACEZR) from the Italian Ministry of Education, University and Research (MIUR) and by PRIN 2022 PNRR from the Italian Ministry of Education, University and Research (MIUR), funded by the European Union - NextGenerationEU - mission 4, component C2, investment 1.1 - P20228M8ZN - CUP F53D23008180001. We thank Gloria Gonzalez Fortes for her contribution in the initial stages of this project

## References

1. N. G. Jablonski, G. Chaplin, The evolution of human skin coloration. J Hum Evol 39, 57– 106 (2000).

2. N. G. Jablonski, The evolution of human skin pigmentation involved the interactions of genetic, environmental, and cultural variables. Pigment Cell Melanoma Res 34, 707–729 (2021).

3. A. R. Rogers, D. Iltis, S. Wooding, Genetic Variation at the MC1R Locus and the Time since Loss of Human Body Hair. Curr Anthropol 45, 105–108 (2004).

4. G. Chaplin, Geographic distribution of environmental factors influencing human skin coloration. Am J Phys Anthropol 125, 292–302 (2004).

5. R. A. Sturm, M. Larsson, Genetics of human iris colour and patterns. Pigment Cell Melanoma Res 22, 544–562 (2009).

6. J. Liu, H. K. Bitsue, Z. Yang, Skin colour: A window into human phenotypic evolution and environmental adaptation. Mol Ecol 33 (2024).

7. L. Chaitanya, et al., The HIrisPlex-S system for eye, hair and skin colour prediction from DNA: Introduction and forensic developmental validation. Forensic Sci Int Genet 35, 123– 135 (2018).

8. S. Walsh, et al., Global skin colour prediction from DNA. Hum Genet 136, 847–863 (2017).

9. S. Walsh, et al., Developmental validation of the HIrisPlex system: DNA-based eye and hair colour prediction for forensic and anthropological usage. Forensic Sci Int Genet 9, 150–161 (2014).

10. M. Ferrando-Bernal, C. M. Brand, J. A. Capra, Inferring human phenotypes using ancient DNA: from molecules to populations. Curr Opin Genet Dev 90, 102283 (2025).

11. J. Draus-Barini, et al., Bona fide colour: DNA prediction of human eye and hair colour from ancient and contemporary skeletal remains. Investig Genet 4, 3 (2013).

12. T. E. King, et al., Identification of the remains of King Richard III. Nat Commun 5, 5631 (2014).

13. S. Brace, et al., Ancient genomes indicate population replacement in Early Neolithic Britain. Nat Ecol Evol 3, 765–771 (2019).

14. I. Olalde, et al., The Beaker phenomenon and the genomic transformation of northwest Europe. Nature 555, 190–196 (2018).

15. T. Z. T. Jensen, et al., A 5700 year-old human genome and oral microbiome from chewed birch pitch. Nat Commun 10, 5520 (2019).

16. M. A. DePristo, et al., A framework for variation discovery and genotyping using next-generation DNA sequencing data. Nat Genet 43, 491–498 (2011).

17. S. Rubinacci, D. M. Ribeiro, R. J. Hofmeister, O. Delaneau, Efficient phasing and imputation of low-coverage sequencing data using large reference panels. Nat Genet 53, 120–126 (2021).

18. R. Nielsen, J. S. Paul, A. Albrechtsen, Y. S. Song, Genotype and SNP calling from nextgeneration sequencing data. Nat Rev Genet 12, 443–451 (2011).

19. R Core Team, R: A Language and Environment for Statistical Computing. R Foundation for Statistical Computing, Vienna, Austria. [Preprint] (2021). Available at: https://www.R-project.org/.

20. Q. Fu, et al., Genome sequence of a 45,000-year-old modern human from western Siberia. Nature 514, 445–449 (2014).

21. T. Günther, et al., Population genomics of Mesolithic Scandinavia: Investigating early post-glacial migration routes and high-latitude adaptation. PLoS Biol 16, e2003703 (2018).

22. A. Seguin-Orlando, et al., Genomic structure in Europeans dating back at least 36,200 years. Science (1979) 346, 1113–1118 (2014).

23. M. E. Allentoft, et al., 100 ancient genomes show repeated population turnovers in Neolithic Denmark. Nature 625, 329–337 (2024).

24. H. Malmström, et al., The genomic ancestry of the Scandinavian Battle Axe Culture people and their relation to the broader Corded Ware horizon. Proceedings of the Royal Society B: Biological Sciences 286, 20191528 (2019).

25. C. E. G. Amorim, et al., Understanding 6th-century barbarian social organization and migration through paleogenomics. Nat Commun 9, 3547 (2018).

26. A. Margaryan, et al., Population genomics of the Viking world. Nature 585, 390–396 (2020).

27. M. Sikora, et al., The population history of northeastern Siberia since the Pleistocene. Nature 570, 182–188 (2019).

28. M. L. Antonio, et al., Stable population structure in Europe since the Iron Age, despite high mobility. Elife 13 (2024).

29. N. Marchi, et al., The genomic origins of the world’s first farmers. Cell 185, 1842-1859.e18 (2022).

30. J. Pearson, et al., Mobility and kinship in the world’s first village societies. Proceedings of the National Academy of Sciences 120 (2023).

31. A. J. Ammerman, “The Transition to Early Farming in Europe” in Simulating Transitions to Agriculture in Prehistory, S. Pardo-Gordó Salvador and Bergin, Ed. (Springer International Publishing, 2021), pp. 225–253.

32. P. Bellwood, First Farmers: The Origins of Agricultural Societies, 2nd Ed. (Wiley-Blackwell, 2023).

33. Y. Kuijpers, et al., Evolutionary Trajectories of Complex Traits in European Populations of Modern Humans. Front Genet 13 (2022).

34. P. Sulem, et al., Genetic determinants of hair, eye and skin pigmentation in Europeans. Nat Genet 39, 1443–1452 (2007).

35. J. Liu, G.C. Black, S.J. Kimber, P.I. Sergouniotis, Generation of a human induced pluripotent stem cell line carrying the TYR c.575C>A (p.Ser192Tyr) and c.1205G>A (p.Arg402Gln) variants in homozygous state using CRISPR-Cas9 genome editing. Stem Cell Res 64, 102880 (2022).

36. R. L. Lamason, et al., SLC24A5, a Putative Cation Exchanger, Affects Pigmentation in Zebrafish and Humans. Science (1979) 310, 1782–1786 (2005).

37. C. Basu Mallick, et al., The Light Skin Allele of SLC24A5 in South Asians and Europeans Shares Identity by Descent. PLoS Genet 9, e1003912 (2013).

38. M. Lin, et al., Rapid evolution of a skin-lightening allele in southern African KhoeSan. Proceedings of the National Academy of Sciences 115, 13324–13329 (2018).

39. L. A. Marano, J. D. Andersen, F. T. Goncalves, A. L. O. Garcia, C. Fridman, Evaluation of HIrisplex-S system markers for eye, skin and hair color prediction in an admixed Brazilian population. Forensic Sci Int Genet Suppl Ser 7, 427–428 (2019).

40. D. M. Hohl, et al., Applicability of the IrisPlex system for eye color prediction in an admixed population from Argentina. Ann Hum Genet 86, 297–327 (2022).

41. T. M. T. Carratto, et al., Evaluation of the HIrisPlex-S system in a Brazilian population sample. Forensic Sci Int Genet Suppl Ser 7, 794–796 (2019).

42. H. Li, et al., The Sequence Alignment/Map format and SAMtools. Bioinformatics 25, 2078– 2079 (2009).

43. A. McKenna, et al., The Genome Analysis Toolkit: A MapReduce framework for analyzing next-generation DNA sequencing data. Genome Res 20, 1297–1303 (2010).

