## Supplementary text for "Inference of human pigmentation from ancient DNA by genotype likelihood"

### Supporting Information Text

#### Materials and Methods

**Conceptual framework.** The HIrisPlex-S system can predict eye, hair, and skin color by analyzing 41 informative genetic variants (1–3). Originally developed for forensic applications, the standard protocol has been recently applied to ancient DNA studies. However, to date, no systematic evaluation has assessed the robustness of this method when working with low-coverage data.
In this study, we quantify the impact of coverage on the inference of the phenotypic traits, following three different approaches described below.
The first protocol follows the classical procedure of HIrisPlex-S (1–3), based on direct genotype calling, called hereby *direct* approach. In the second protocol, the phenotypic inference is based on imputed genomes, following the pipeline specifically developed for ancient DNA data (4), called hereby *imputation* approach. The third protocol makes use of a *probabilistic* approach based on the computation of genotype likelihood (5) to explicitly take into account the uncertainty linked to low-coverage data in the characterization of genomic variants.
All the three approaches have been tested using two ancient high-coverage genomes, which were downsampled considering different coverage levels. The outcomes of this validation step helped to identify the coverage threshold at which each approach should be applied to achieve the most reliable phenotypic predictions. Lastly, we applied these approaches to a broader ancient genomic dataset described below.

**Genomic dataset.** The ancient genomic dataset comprises 348 samples from the Upper Paleolithic to the Iron Age, encompassing the geographic region of Eurasia, collected from the literature (S1_Appendix). We selected whole-genome sequences with an average coverage of at least 1x.
For the validation step, we selected two high-coverage samples suitable for downsampling process. The main criteria were to ensure the representation of different historical periods and to select those with the highest coverage at the 41 HIrisPlex-S positions. Based on these, we selected two individuals from the Paleolithic and Mesolithic periods:

- Ust'-Ishim, 45,045 calBP from western Siberia (6);
- SF12, 9,033±8,757 calBP from Sweden (7).

**Bioinformatics analysis.** All the samples were processed following the guidelines specifically designed for ancient genome analysis (8). For Ust'-Ishim and SF12 samples, the FastQ files were downloaded from *ENA* and analyzed with *FastQC* v0.12.1 (9) for quality control of the DNA fragments and adapter sequences were removed using *AdapterRemoval* v2.3.1 (10). The filtered reads were then aligned using *BWA* software v0.7.17 246 (11) with the *aln* algorithm. The minimum read length (*--minlength 30*), base quality (*--minquality 30*) and mapping quality (*-q 30*) filters were applied, and seeding was disabled (*-l 1000*) to ensure alignment of all reads against the GRCh37.p13/hg19 human reference genome (12). The final autosomal average coverage was calculated using *SAMtools depth* command v1.11 (13) with the option *-a*, to include all positions, even those with 0 depth.

Then, we generated a *pileup* file for each sample, containing the alignment information for the 41 HIrisPlex-S positions, using *SAMtools mpileup* command (v1.11) (13). The indel at chromosome 16 position 89985753 (rs312262906, insertion of an "A" nucleotide) was assessed using *VarScan* v2.3.9 (14) algorithm *mpileup2indel* from the *SAMtools mpileup* output file. The presence of the indel was confirmed if "=+A" was reported at chromosome 16 positions 89985750, 89985751, 89985752, 89985753 or 89985754.

**Testing the robustness of phenotypic inference on ancient data.** The principle underlying the downsampling procedure is to accurately evaluate the amount of information that might be lost at lower coverage levels by artificially reducing the sequencing depth (i.e., the number of times a single nucleotide is sequenced). We followed two different downsampling procedures: 1) a pointwise progressive downsampling applied to the validation of the *direct* and *probabilistic* approaches, and 2) mean progressive downsampling applied to the validation of the *imputation* approach.

To achieve incremental pointwise downsampling at each stage, the *SAMtools pileup* file was employed to detect read identifiers ("*--output-extra QNAME*" flag) and then, using an *in-house* R script (15), we employed a non-replacement sampling to ensure random selection of unique elements to prevent read and nucleotide duplication (using the function *sample* with the option *replace = false*). Subsequently, we extracted reads the original BAM files from chromosomes 5, 6, 9, 11, 12, 14, 15, and 20, using the sampled identifiers, with *SAMtools* (v1.11) *view* command (13), resulting in a single BAM file per chromosome. Chromosome 16 underwent a similar process but required 14 separate BAM files, one for each genomic position, to manage read overlap and potential coverage increases due to the proximity of the positions them-selves. For the mean progressive downsampling, we used *SAMtools* (v1.11) *view* command (13) with the *-s* flag to output a subset of the input alignment in order to achieve the target coverage levels. The process began with the high-coverage genome, progressively reducing it to 1x coverage. To ensure randomness in read selection, distinct seed values were employed for each downsampling operation, conducted on separate computing nodes. We validated the accuracy of the achieved coverage using the *SAMtools* v1.11 *depth -a* command (13).

- **Direct approach.** Variants were called on BAM files using the *GATK* (Genome Analysis Toolkit) *UnifiedGenotyper* algorithm v3.5 (16). Key parameters included a minimum confidence threshold for calling variants (*-stand_call_conf* = 30), a minimum confidence threshold for emitting variants (*-stand_emit_conf* = 20) and a minimum base quality threshold (*-mbq* = 30). The analysis was configured to output all sites (*-out_mode EMIT_ALL_SITES*), regardless of whether they contain variant or invariant alleles compared to the reference genome (GRCh37.p13). Additionally, the *-glm BOTH* option was employed to simultaneously compute genotype likelihoods using both SNP and indel models. The option *-L* was used to compute calling only on HIrisPlex-S sites.

- **Imputation approach.** We tested the aDNA imputation pipeline proposed in Sousa da Mota *et al.* (2023) to evaluate how phenotypic predictions change with different average coverage levels and the presence of missing variants. All the steps were performed on samples subjected to mean progressive downsampling. Following the guidelines of Sousa da Mota, the reference panel selected was the 1000 Genomes phase 3 v5, comprising 3,202 samples re-sequenced at 30x (4, 17). Data were downloaded in VCF format from NCBI (http://ftp.1000genomes.ebi.ac.uk/vol1/ftp/data_collections/1000G_2504_high_coverage/). Since the ancient samples comprised in our database were aligned against the GRCh37.p13 reference genome and the reference panel was aligned to the GRCh38 reference genome, we performed the liftOver of the reference panel using *Picard liftoverVCF* v2.24.1 (18) with the *hg38ToHg19* chain from the University of California, Santa Cruz liftOver tool (19). *BCFtools view* command (v1.11) (20) was used with the options *-S* and *-m 2 -M 2 -v snps* to remove related individuals and multiallelic records, resulting in approximately 90 million SNPs and 2,504 individuals. Before proceeding with the imputation step, the HIrisPlex-S positions were masked from all the mean progressive downsampled genomes. The imputation of the downsampled genomes was performed using *GLIMPSE* v1.1.1 (21). Chromosomes were divided into chunks of sizes in the range 1–2 Mb using *GLIMPSE_chunk*, with a 200-kb buffer region at each side of a chunk. Imputation was then carried out with *GLIMPSE_phase* using parameters *--burn 10*, *--main 15* and *--pbwt-depth2*. Finally, the imputed chunks were ligated using *GLIMPSE_ligate* and the most likely haplotype pair for each sample was determined using the *--solve* option in *GLIMPSE_sample*. The final output was a VCF file containing all the called and phased genotypes. Of the 41 HIrisPlex-S positions, only 39 were successfully retrieved due to the absence of the variants (rs312262906 and rs201326893) in the reference panel. The extracted HIrisPlex-S genotypes were then used for phenotypic predictions.

- **Probabilistic approach.** The standard protocol of the HIrisPlex-S system allows a single phenotypic prediction per sample by taking as input an informative allele from a single genotype call for each of the 41 positions.

  Two substantial modifications have been implemented in the *probabilistic* approach proposed here: 1) the variant calling phase, where instead of exclusively adopting the most probable genotype proposed by *GATK*, all the 10 possible genotypes are considered based on their respective genotype likelihoods (5). This approach ensures that multiple genotypes may be evaluated for the same position according to their genotype likelihood values, facilitating a more comprehensive analysis; 2) the number of predictions made for each sample.

We conducted, indeed, 1,000 samplings from the 10 possible genotypes, based on their genotype likelihoods and posterior probabilities. Each sampling generated a distinct combination of informative alleles. This transition from one prediction per sample to 1,000 predictions per sample, encompasses all possible genotype combinations across the 41 positions, acknowledging the inferential uncertainty linked to low coverage data.

All the steps were performed starting from the *SAMtools pileup* file and all the computations were performed using logarithmic values to avoid machine underflow. Genotype likelihoods for all the ten possible genotypes (given the assumption of a diploid individual, i.e., AA, AC, AG, AT, CC, CG, CT, GG, GT and TT) were calculated following the formula of the first version of *GATK* (dragon) (22):

$$Pr(D|G = \{A_1, A_2\}) = \prod_{i=1}^{M} Pr(b_i|G = \{A_1, A_2\}) = \prod_{i=1}^{M} \left(\frac{1}{2}Pr(b_i|A_1) + \frac{1}{2}Pr(b_i|A_2)\right)$$

$$Pr(b|A) = \begin{cases} \frac{e}{3} : b \neq A \\ 1 - e : b = A \end{cases}$$

considering the base read during sequencing ($b_i$), its respective depth of coverage ($M$) and the probability of sequencing error, calculated from the Phred scaled quality score $e = 10^{-q/10}$.

In addition to computing the genotype likelihood, a prior probability for each genotype must be defined to obtain posterior probabilities for all the genotypes (5). The prior genotype probability chosen was the same used by *GATK UnifiedGenotyper* v3.5, i.e. the expected probability that an individual is heterozygous at a given locus. For humans, a default heterozygosity value (hets) of 0.001 is set (16).

Lastly, genotypes were sampled using the *sample* function (*R* environment v4.3.3 (15)), considering their genotype posterior probabilities. The likelihood of selecting a particular genotype for sampling thus depended on the strength of support it received from the sequencing data, with genotypes possessing higher posterior probabilities being more likely to be sampled.

**Phenotypic prediction.** The set of 41 SNPs required by the HIrisPlex-S system suffers from a technical issue, the strand definition. For each *pileup* file we checked the strand for each of the 41 SNPs by comparing the reference and alternate alleles in our dataset with the informative allele from the HIrisPlex-S system. The final result consists of a table containing one or 1,000 rows (depending on the methodological approach chosen), and 41 columns, one for each informative allele. The information present in the columns needs to be in the format required by the HIrisPlex-S website:

- The absence of the input allele is defined by “0”;

- An heterozygous condition (one informative allele present) is defined by “1”;
- A homozygous condition (two copies of the informative allele present) is defined by “2”;
- A missing SNP is defined by “NA”.

This table was saved as a *CSV* file, formatted for compatibility with the HIrisPlex-S website as specified in the HIrisPlex-S Webtool User Manual Version 2.0 (2018 (1–3)). The prediction results obtained for each sample were as a *CSV* file and then analyzed in *R* environment v4.3.3 (15) using an *in-house* R script, following the guidelines outlined in the HIrisPlex-S user Webtool User Manual Version 2.0 (2018) (1–3), to obtain the final phenotypic traits inferences (Figures S1-S6).

Moreover, when applying the *probabilistic* approach, phenotypic predictions meeting a 90% confidence level (i.e., the same phenotype predicted in at least 900 out of 1,000 iterations) were considered reliable and reported as unique (Figures S7-S12). This threshold minimizes errors arising from stochastic associations between genotypes with lower posterior probabilities, ensuring that only highly confident predictions are included.

For the phenotypic inference of Eurasian samples, predictions below the 90% threshold were classified as uncertain and excluded from the final interpretation, as they did not meet the required accuracy.

To improve the clarity of results, we grouped phenotypic predictions into macro-groups by color intensity:

- Eye pigmentation: three categories (Brown estimates as Dark; Intermediate estimates as Intermediate; Blue estimates as Light).
- Hair pigmentation: four categories (Black, Dark brown/black, and Brown/dark brown estimates as Dark; Brown and Dark blond estimates as Intermediate; Blond estimates as Light; Red estimates as Red).
- Skin pigmentation: three categories (Dark to black and Dark estimates as Dark; Intermediate estimates as Intermediate; Pale and Very Pale estimates as Light).

This categorization addressed the challenge of differentiating closely related shades, as well as the difficulties observed when working with admixed populations, as highlighted by Carratto *et al.* 2015, Marano *et al.*, 2020 and Hohl *et al.*, 2022 (23–25) (Figures S13-S24; Appendix files from S2 to S6).

### Figure S1

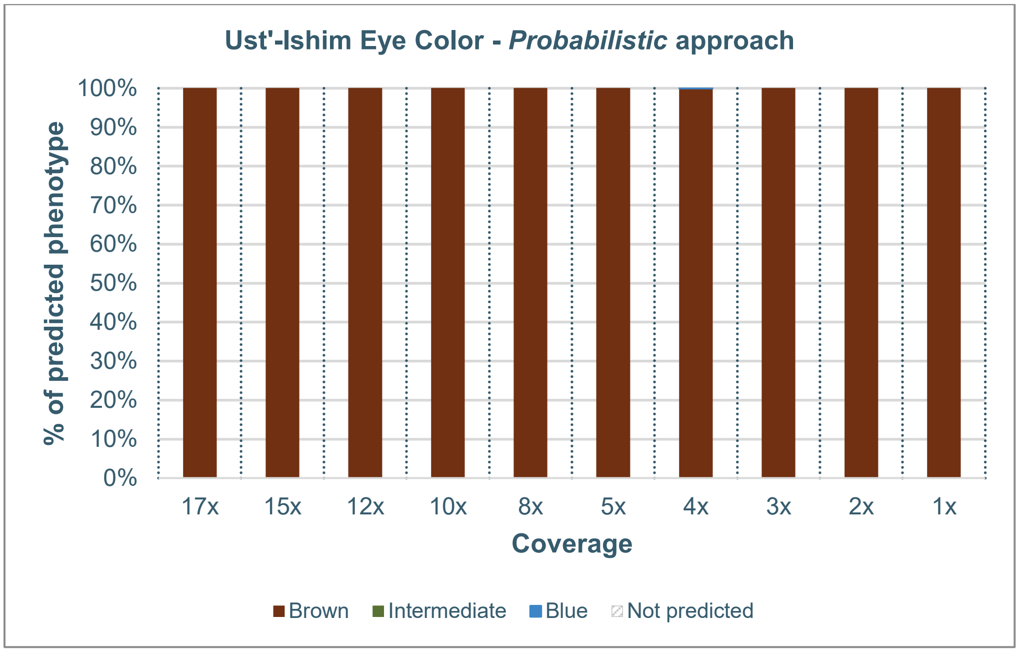


***Eye color phenotypic predictions for the Ust'-Ishim sample obtained through the probabilistic approach at varying sequencing coverages, displaying all prediction outcomes.*** *The x-axis indicates sequencing coverage, while the y-axis represents the percentage of predicted outcomes based on 1000 predictions repeated across 10 iterations.*

### Figure S2

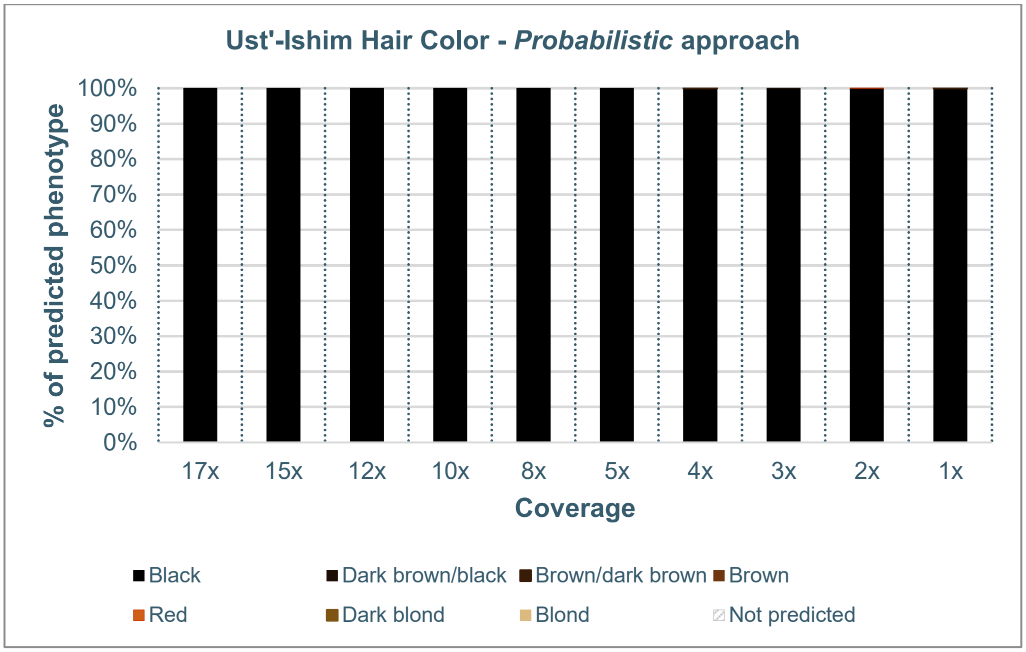


***Hair color phenotypic predictions for the Ust'-Ishim sample obtained through the probabilistic approach at varying sequencing coverages, displaying all prediction outcomes.*** *The x-axis indicates sequencing coverage, while the y-axis represents the percentage of predicted outcomes based on 1000 predictions repeated across 10 iterations.*

### Figure S3

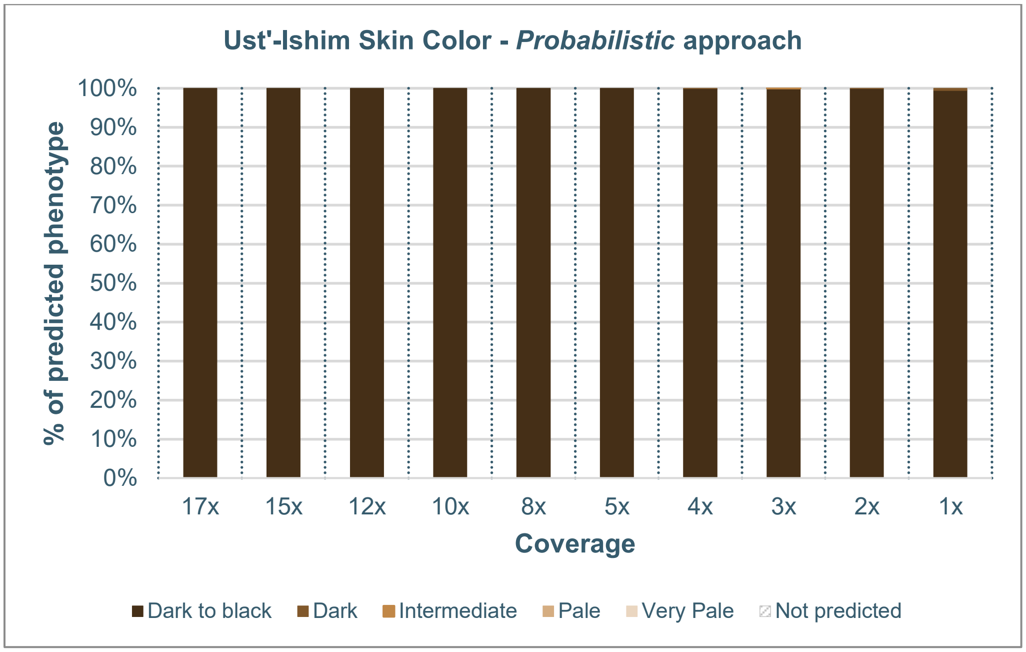


***Skin color phenotypic predictions for the Ust'-Ishim sample obtained through the probabilistic approach at varying sequencing coverages, displaying all prediction outcomes.*** *The x-axis indicates sequencing coverage, while the y-axis represents the percentage of predicted outcomes based on 1000 predictions repeated across 10 iterations.*

### Figure S4

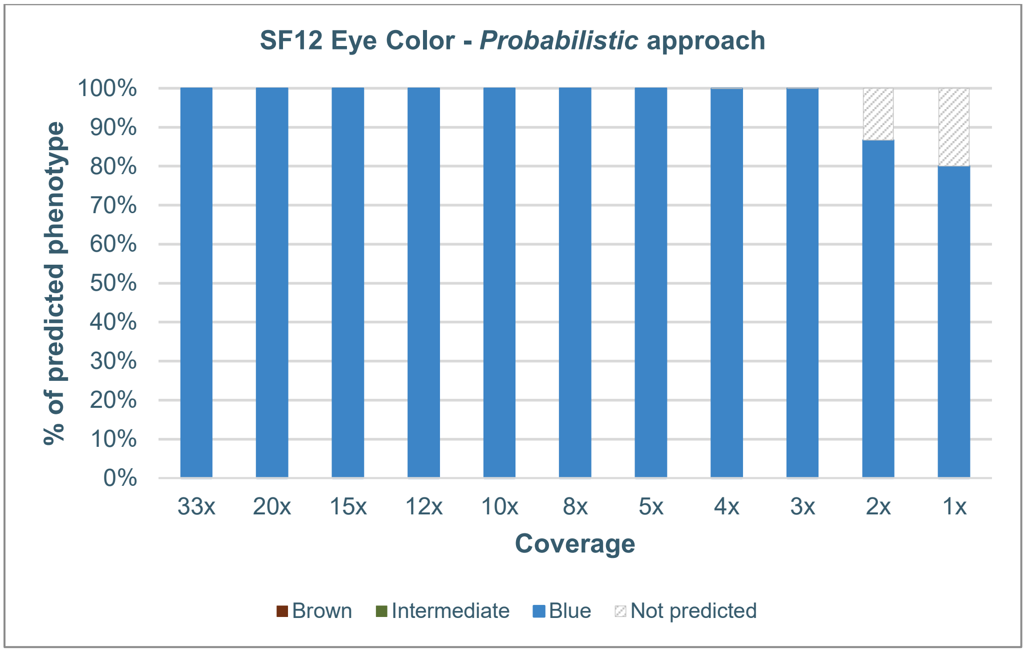


***Eye color phenotypic predictions for the SF12 sample obtained through the probabilistic approach at varying sequencing coverages, displaying all prediction outcomes.*** *The x-axis indicates sequencing coverage, while the y-axis represents the percentage of predicted outcomes based on 1000 predictions repeated across 10 iterations.*

### Figure S5

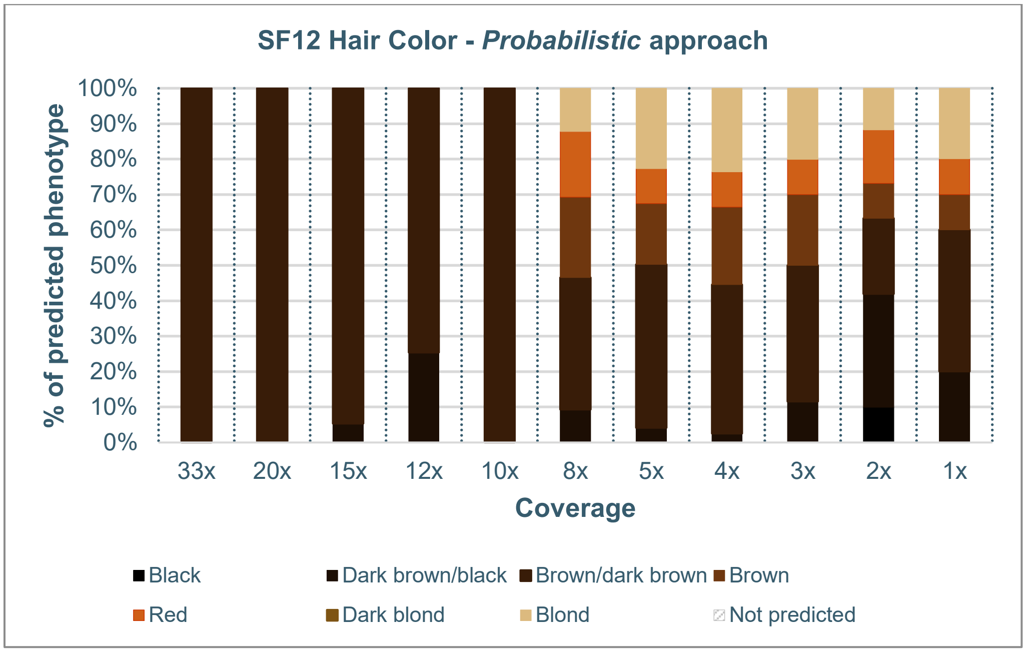


***Hair color phenotypic predictions for the SF12 sample obtained through the probabilistic approach at varying sequencing coverages, displaying all prediction outcomes.*** *The x-axis indicates sequencing coverage, while the y-axis represents the percentage of predicted outcomes based on 1000 predictions repeated across 10 iterations.*

### Figure S6

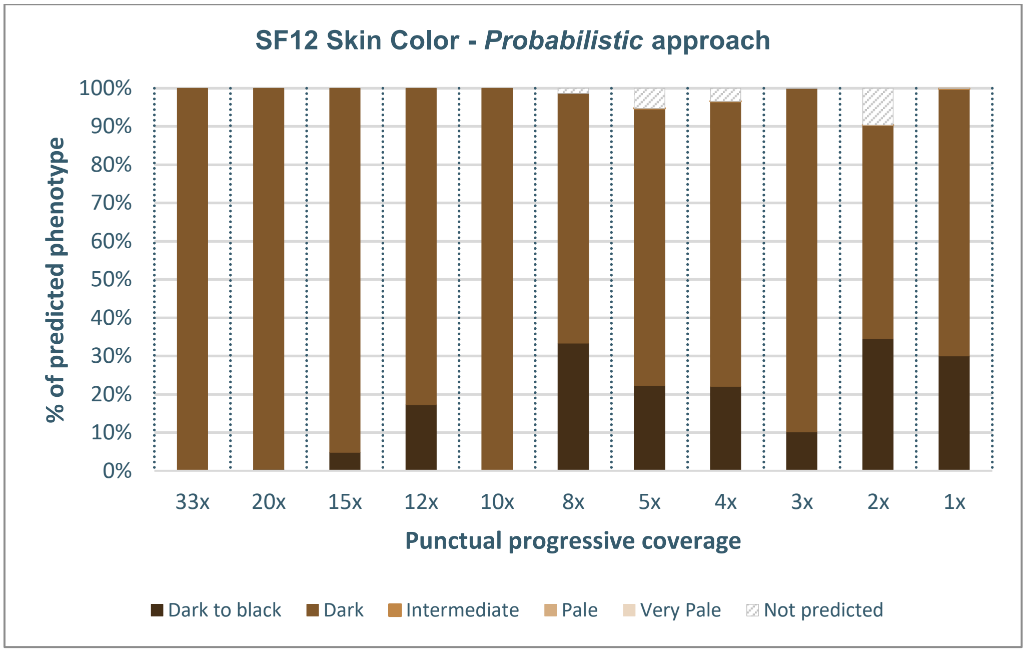


***Skin color phenotypic predictions for the SF12 sample obtained through the probabilistic approach at varying sequencing coverages, displaying all prediction outcomes.*** *The x-axis indicates sequencing coverage, while the y-axis represents the percentage of predicted outcomes based on 1000 predictions repeated across 10 iterations.*

### Figure S7

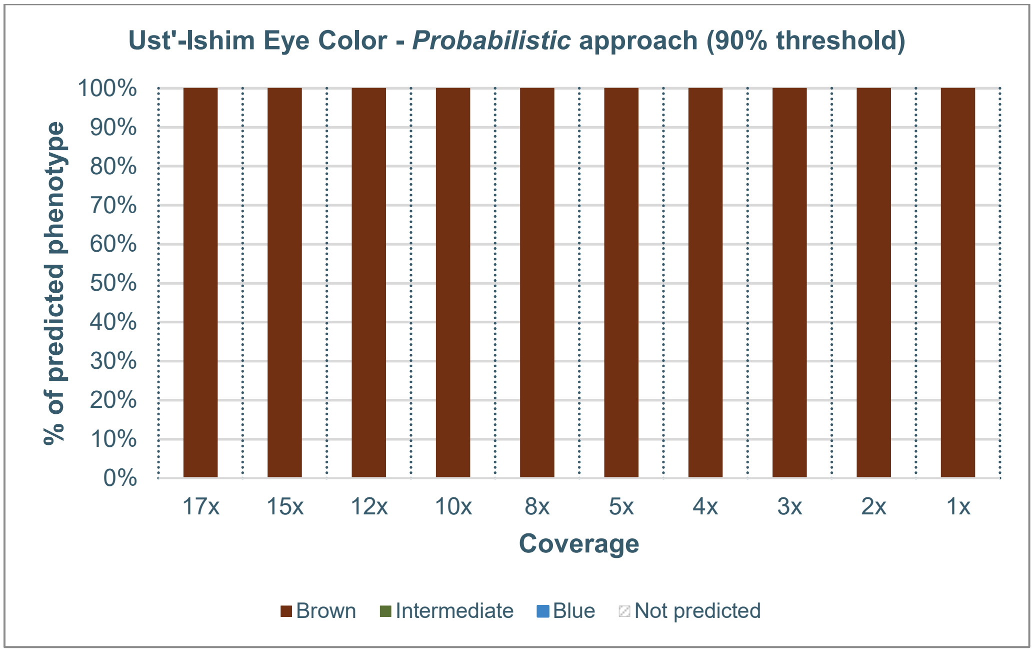


***Eye color phenotypic predictions for the Ust'-Ishim sample obtained through the probabilistic approach at varying sequencing coverages, applying a 90% confidence threshold.** The x-axis indicates sequencing coverage, while the y-axis represents the percentage of predicted outcomes based on 1000 predictions repeated across 10 iterations. We show a unique phenotypic prediction in downsampling experiments where the same phenotype is predicted in at least 900 iterations out of 1,000.*

**Figure S8**

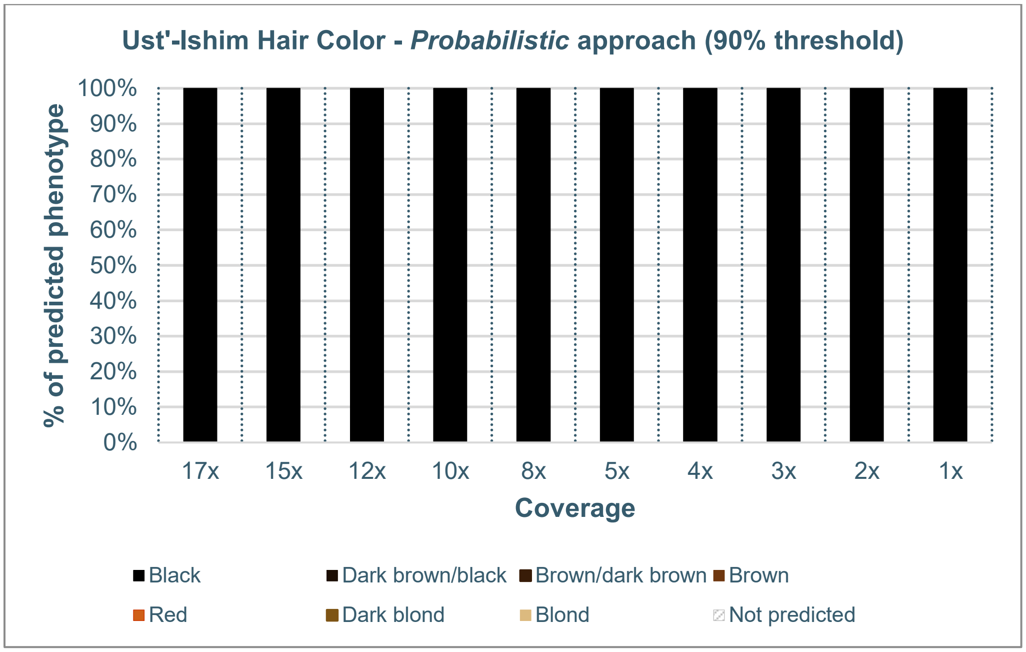


***Hair color phenotypic predictions for the Ust'-Ishim sample obtained through the probabilistic approach at varying sequencing coverages, applying a 90% confidence threshold.*** *The x-axis indicates sequencing coverage, while the y-axis represents the percentage of predicted outcomes based on 1000 predictions repeated across 10 iterations. We show a unique phenotypic prediction in downsampling experiments where the same phenotype is predicted in at least 900 iterations out of 1,000.*

### Figure S9

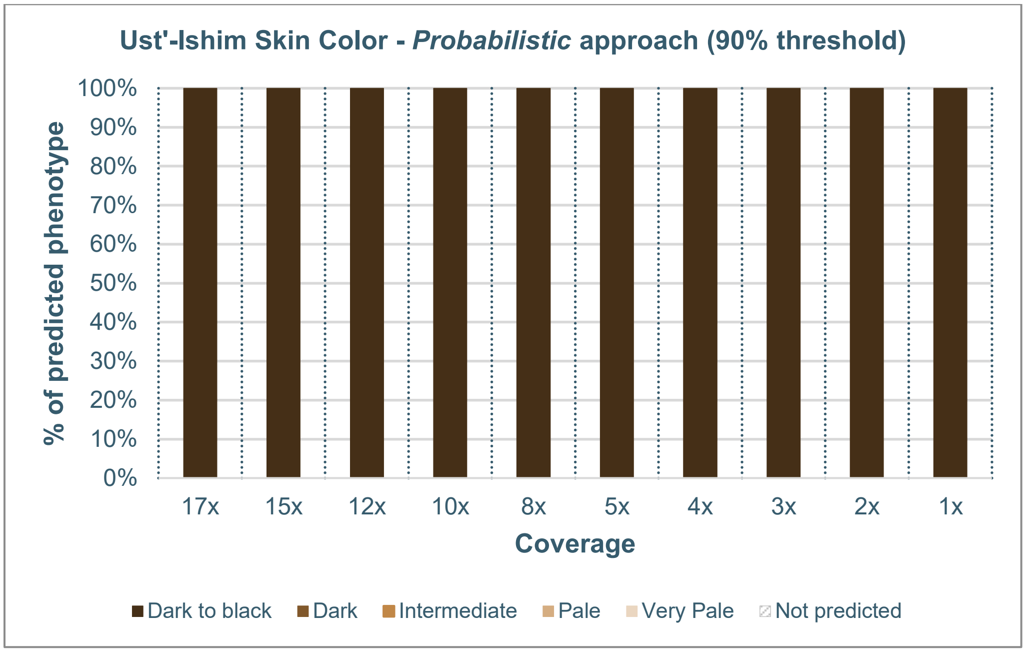


***Skin color phenotypic predictions for the Ust'-Ishim sample obtained through the probabilistic approach at varying sequencing coverages, applying a 90% confidence threshold.*** *The x-axis indicates sequencing coverage, while the y-axis represents the percentage of predicted outcomes based on 1000 predictions repeated across 10 iterations. We show a unique phenotypic prediction in downsampling experiments where the same phenotype is predicted in at least 900 iterations out of 1,000.*

### Figure S10

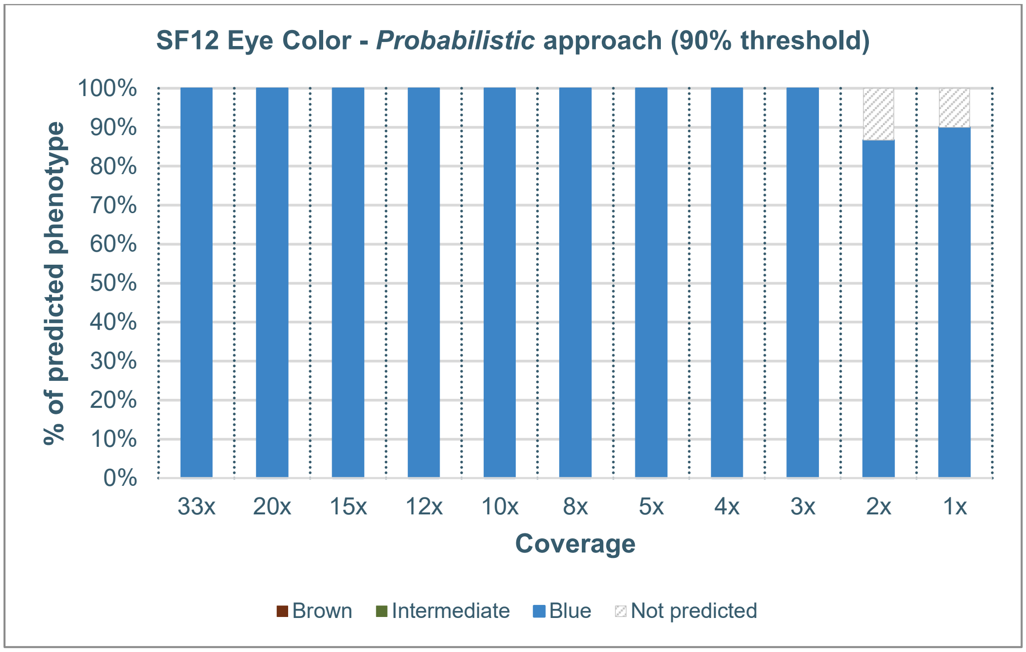


***Eye color phenotypic predictions for the SF12 sample obtained through the probabilistic approach at varying sequencing coverages, applying a 90% confidence threshold.*** *The x-axis indicates sequencing coverage, while the y-axis represents the percentage of predicted outcomes based on 1000 predictions repeated across 10 iterations. We show a unique phenotypic prediction in downsampling experiments where the same phenotype is predicted in at least 900 iterations out of 1,000.*

### Figure S11

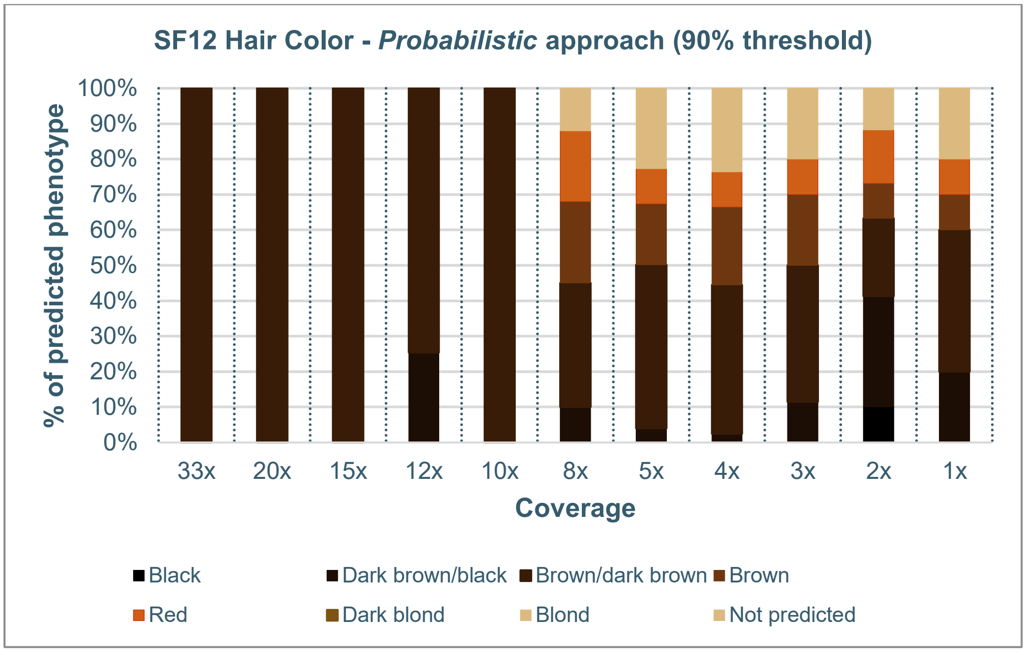


***Hair color phenotypic predictions for the SF12 sample obtained through the probabilistic approach at varying sequencing coverages, applying a 90% confidence threshold.*** *The x-axis indicates sequencing coverage, while the y-axis represents the percentage of predicted outcomes based on 1000 predictions repeated across 10 iterations. We show a unique phenotypic prediction in downsampling experiments where the same phenotype is predicted in at least 900 iterations out of 1,000.*

**Figure S12**

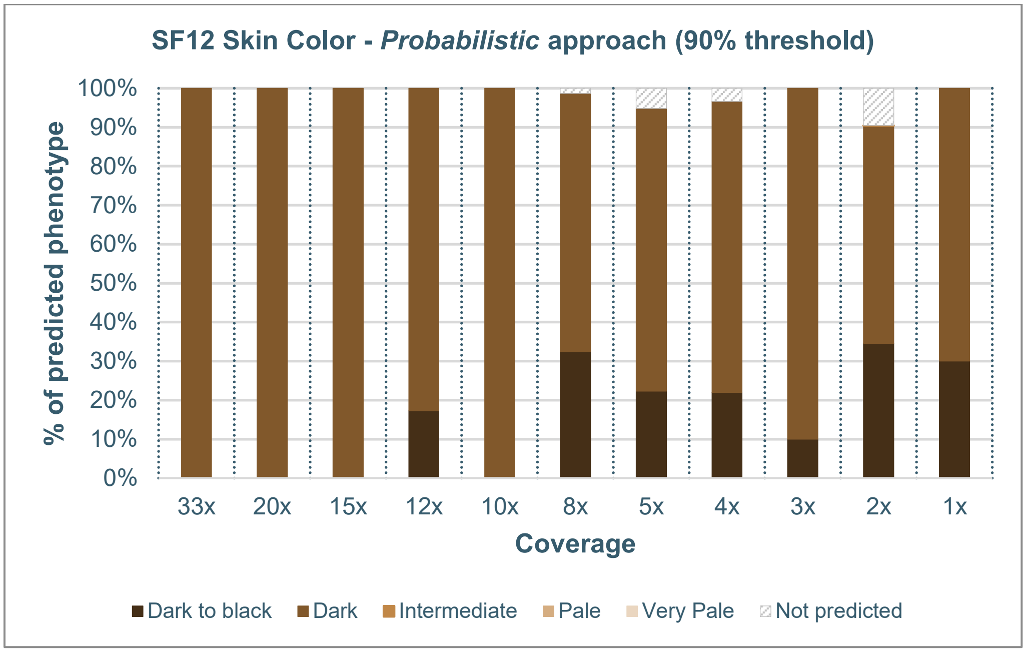


***Skin color phenotypic predictions for the SF12 sample obtained through the probabilistic approach at varying sequencing coverages, applying a 90% confidence threshold.*** *The x-axis indicates sequencing coverage, while the y-axis represents the percentage of predicted outcomes based on 1000 predictions repeated across 10 iterations. We show a unique phenotypic prediction in downsampling experiments where the same phenotype is predicted in at least 900 iterations out of 1,000.*

### Figure S13

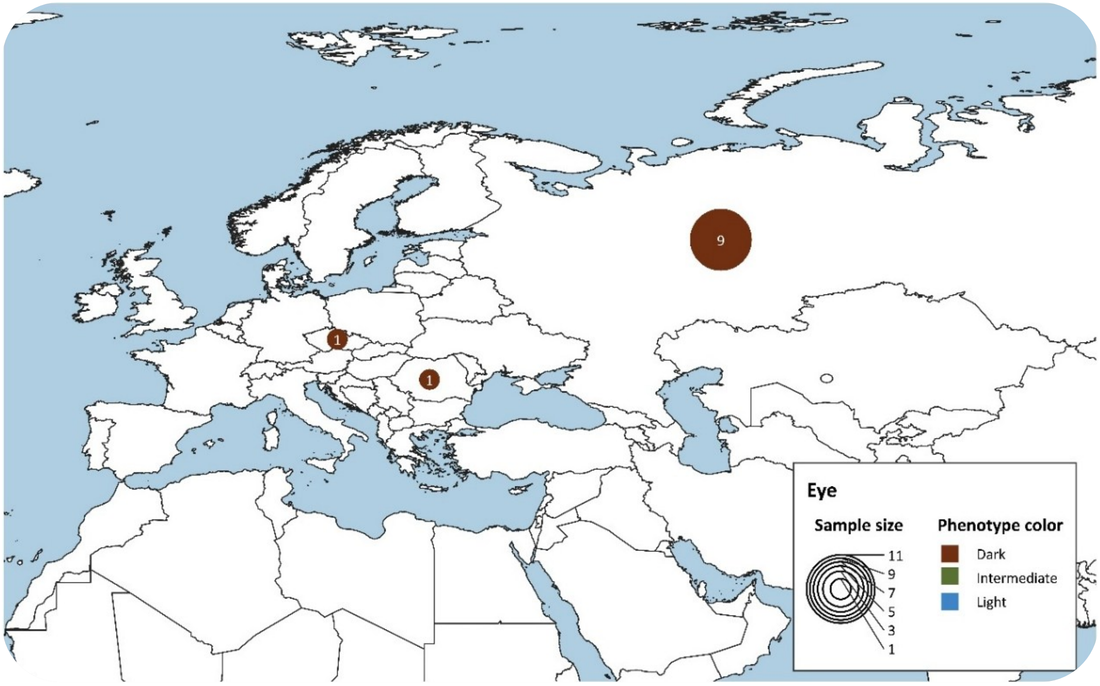


***Temporal and geographical distribution of eye color estimates in Eurasia during the Paleolithic.*** *The map illustrates the spatial distribution of the inferred eye color phenotypes for the Paleolithic period. The size of each pie chart corresponds to the sample size. Eye color results are grouped into 3 categories: Dark (Brown), Intermediate, and Light (Blue).*

### Figure S14

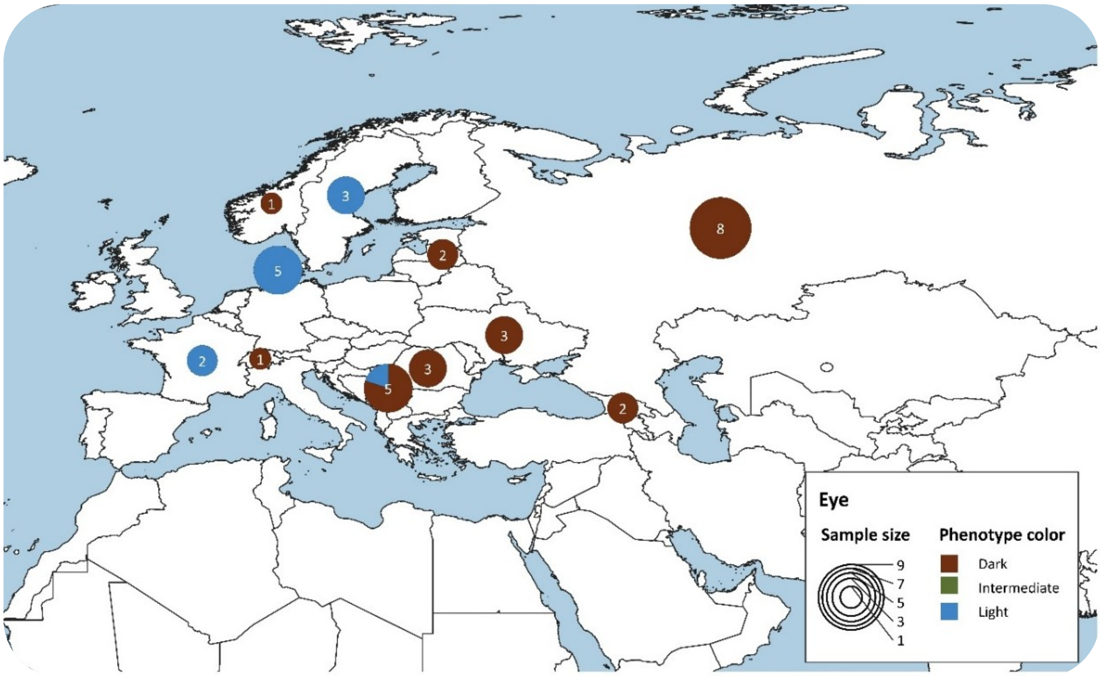


***Temporal and geographical distribution of eye color estimates in Eurasia during the Mesolithic.*** *The map illustrates the spatial distribution of the inferred eye color phenotypes for the Mesolithic period. The size of each pie chart corresponds to the sample size. Eye color results are grouped into 3 categories: Dark (Brown), Intermediate, and Light (Blue).*

### Figure S15

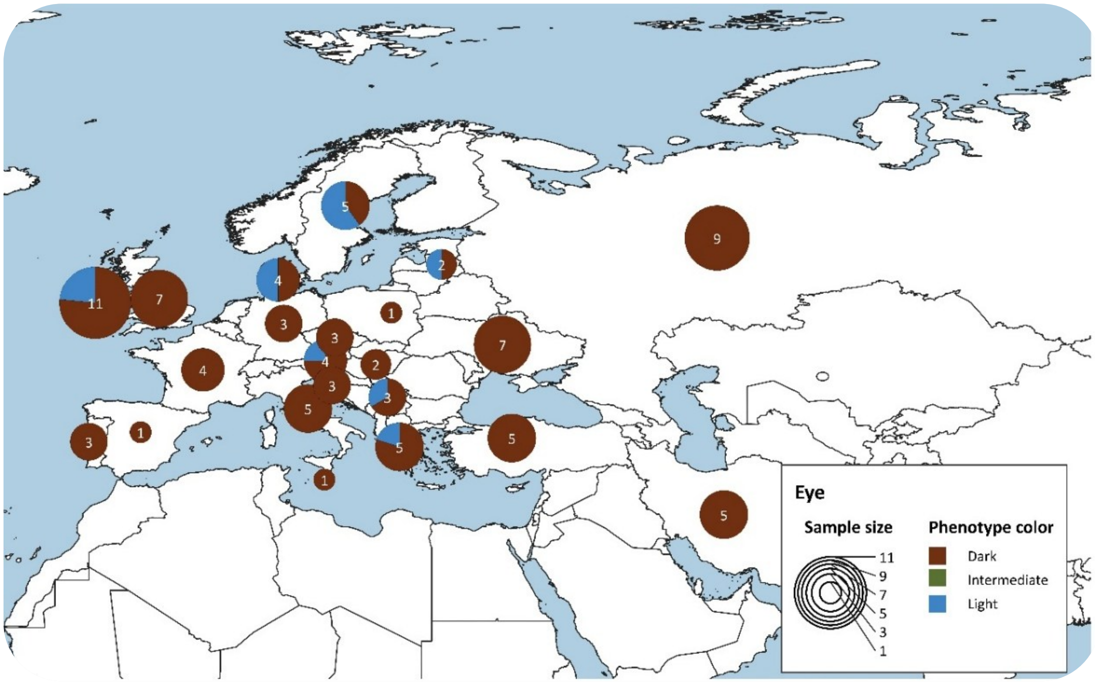


***Temporal and geographical distribution of eye color estimates in Eurasia during the Neolithic.*** *The map illustrates the spatial distribution of the inferred eye color phenotypes for the Neolithic period. The size of each pie chart corresponds to the sample size. Eye color results are grouped into 3 categories: Dark (Brown), Intermediate, and Light (Blue).*

### Figure S16

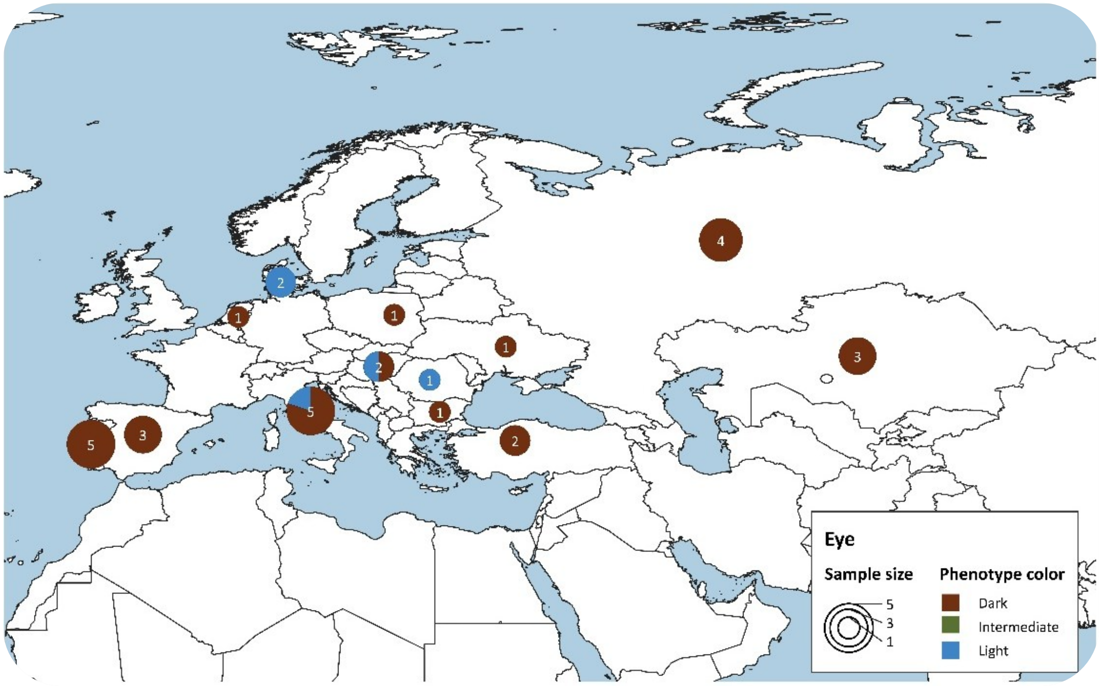


***Temporal and geographical distribution of eye color estimates in Eurasia during the Copper Age.*** *The map illustrates the spatial distribution of the inferred eye color phenotypes for the Copper Age period. The size of each pie chart corresponds to the sample size. Eye color results are grouped into 3 categories: Dark (Brown), Intermediate, and Light (Blue).*

### Figure S17

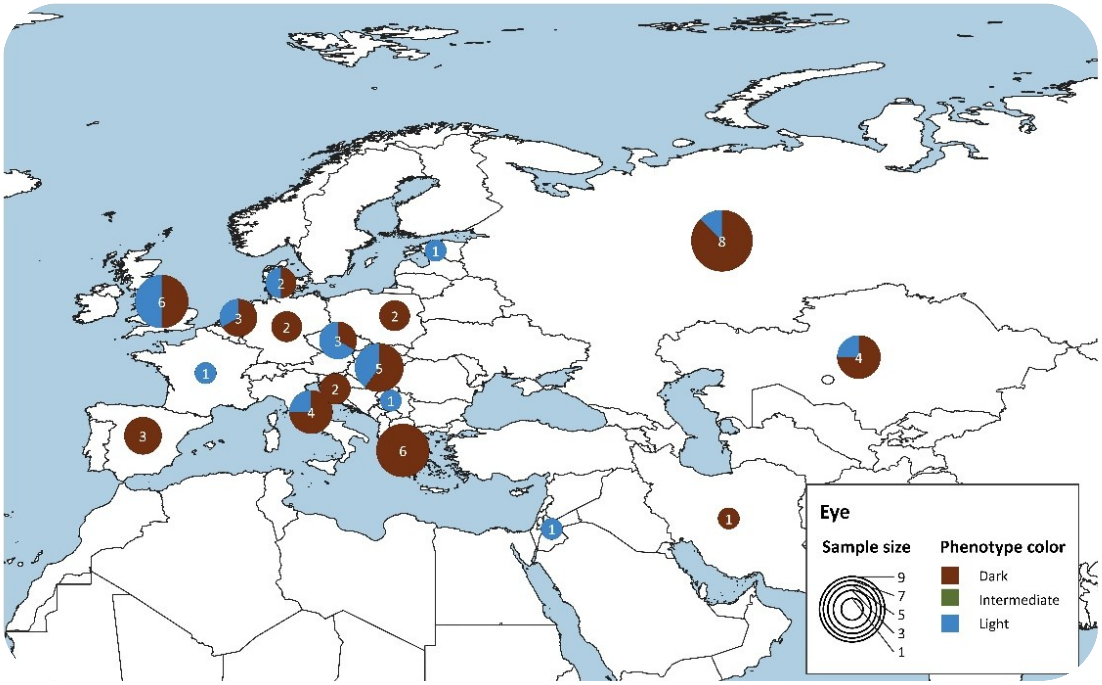


***Temporal and geographical distribution of eye color estimates in Eurasia during the Bronze Age.*** *The map illustrates the spatial distribution of the inferred eye color phenotypes for the Bronze Age period. The size of each pie chart corresponds to the sample size. Eye color results are grouped into 3 categories: Dark (Brown), Intermediate, and Light (Blue).*

### Figure S18

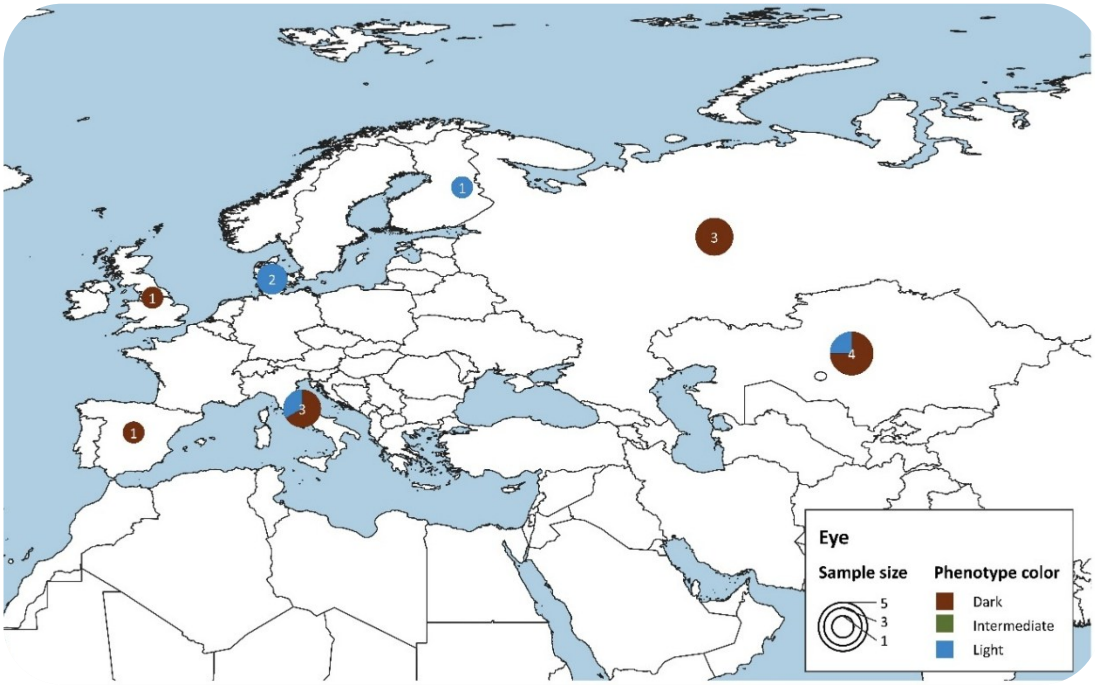


***Temporal and geographical distribution of eye color estimates in Eurasia during the Iron Age.*** *The map illustrates the spatial distribution of the inferred eye color phenotypes for the Iron Age period. The size of each pie chart corresponds to the sample size. Eye color results are grouped into 3 categories: Dark (Brown), Intermediate, and Light (Blue).*

### Figure S19

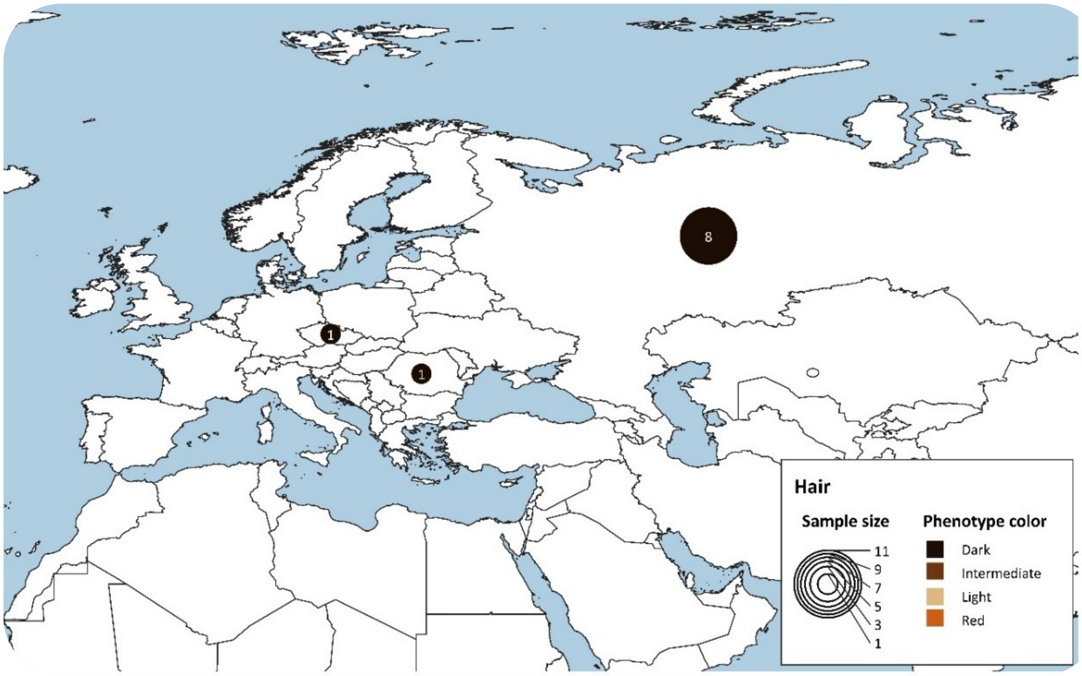


***Temporal and geographical distribution of hair color estimates in Eurasia during the Paleolithic.*** *The map illustrates the spatial distribution of the inferred eye color phenotypes for the Paleolithic period. The size of each pie chart corresponds to the sample size. Eye color results are grouped into 4 categories: Dark (Black, Dark brown/black, and Brown/dark brown estimates), Intermediate (Brown and Dark blond), Light (Blond), and Red.*

### Figure S20

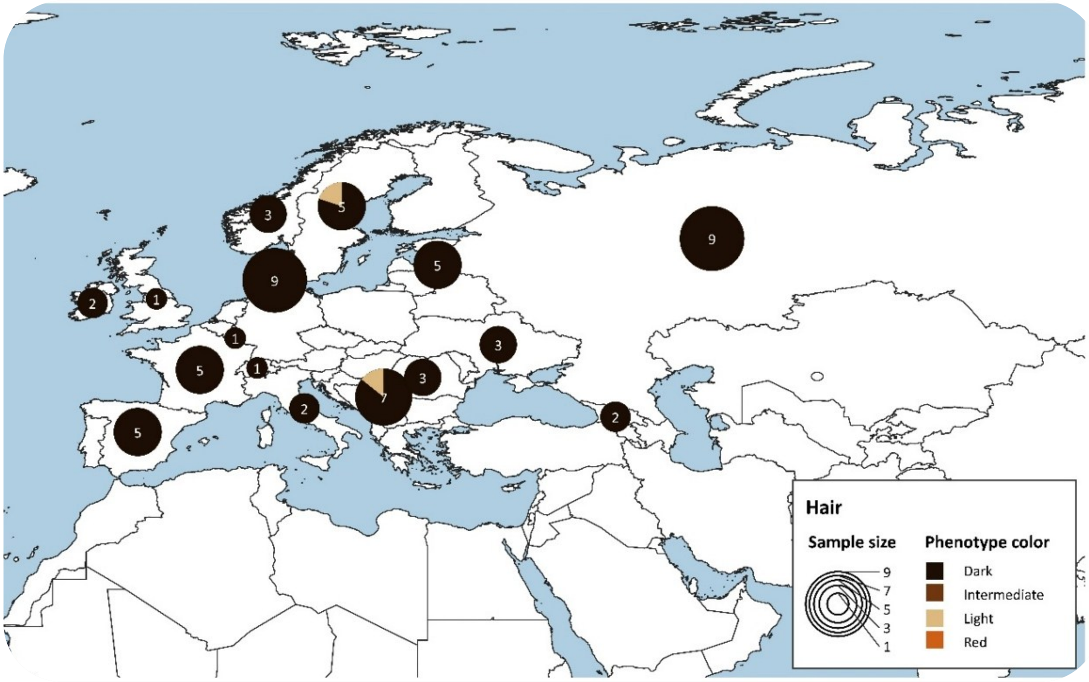


***Temporal and geographical distribution of hair color estimates in Eurasia during the Mesolithic.*** *The map illustrates the spatial distribution of the inferred eye color phenotypes for the Mesolithic period. The size of each pie chart corresponds to the sample size. Eye color results are grouped into 4 categories: Dark (Black, Dark brown/black, and Brown/dark brown estimates), Intermediate (Brown and Dark blond), Light (Blond), and Red.*

### Figure S21

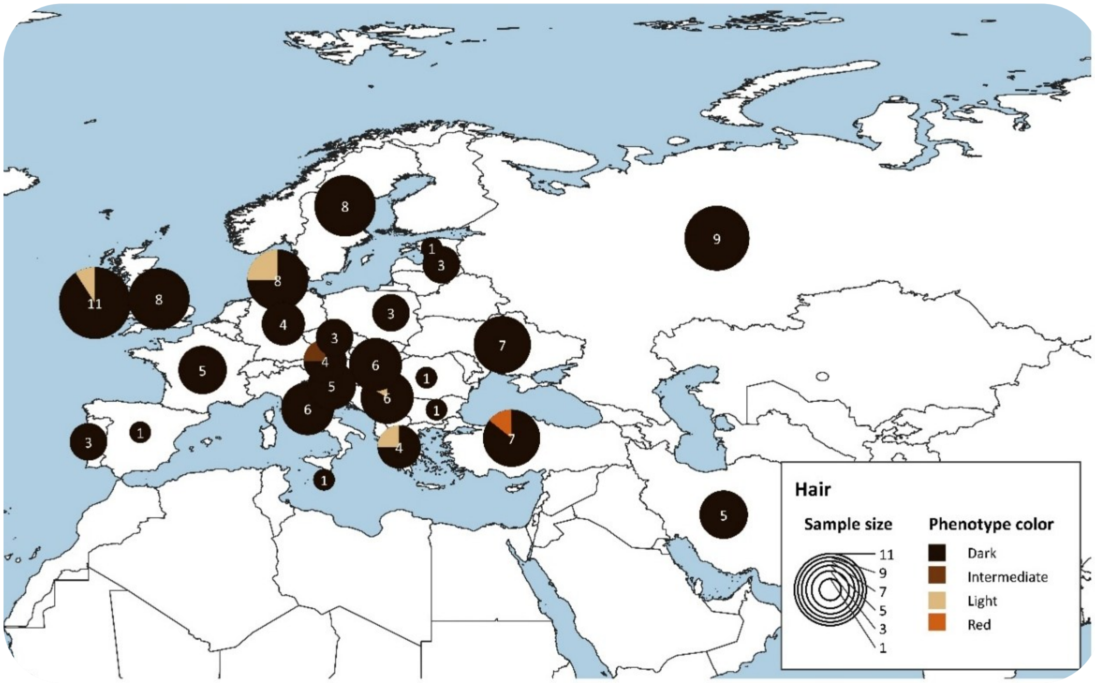


***Temporal and geographical distribution of hair color estimates in Eurasia during the Neolithic.*** *The map illustrates the spatial distribution of the inferred eye color phenotypes for the Neolithic period. The size of each pie chart corresponds to the sample size. Eye color results are grouped into 4 categories: Dark (Black, Dark brown/black, and Brown/dark brown estimates), Intermediate (Brown and Dark blond), Light (Blond), and Red.*

### Figure S22

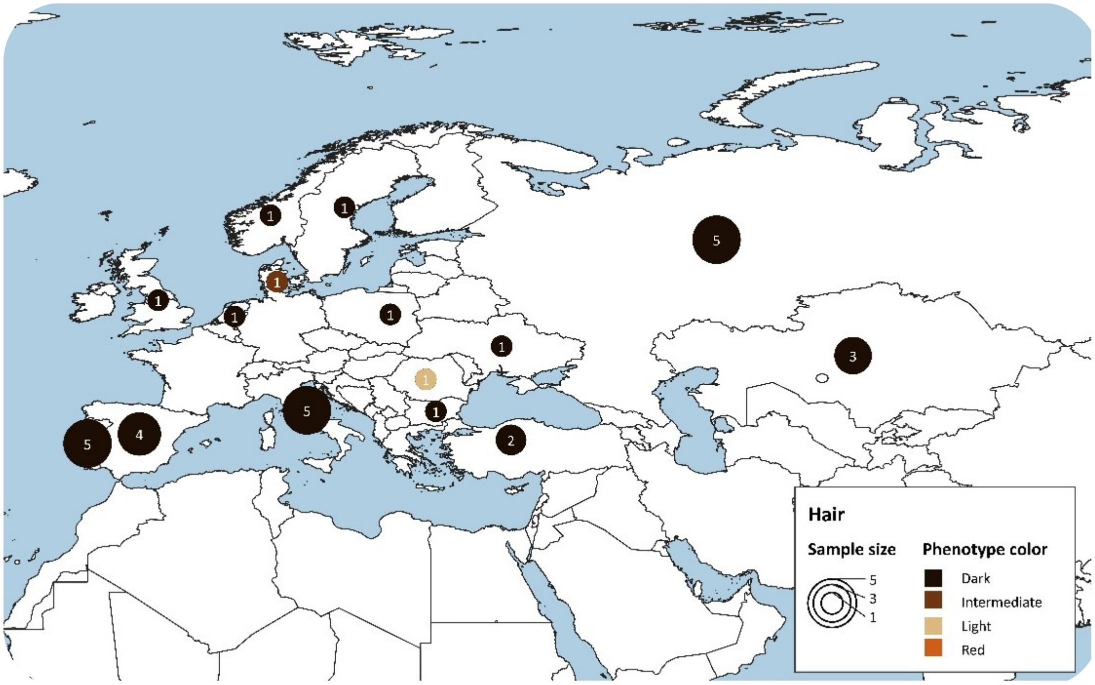


***Temporal and geographical distribution of hair color estimates in Eurasia during the Copper Age.*** *The map illustrates the spatial distribution of the inferred eye color phenotypes for the Copper Age period. The size of each pie chart corresponds to the sample size. Eye color results are grouped into 4 categories: Dark (Black, Dark brown/black, and Brown/dark brown estimates), Intermediate (Brown and Dark blond), Light (Blond), and Red.*

### Figure S23

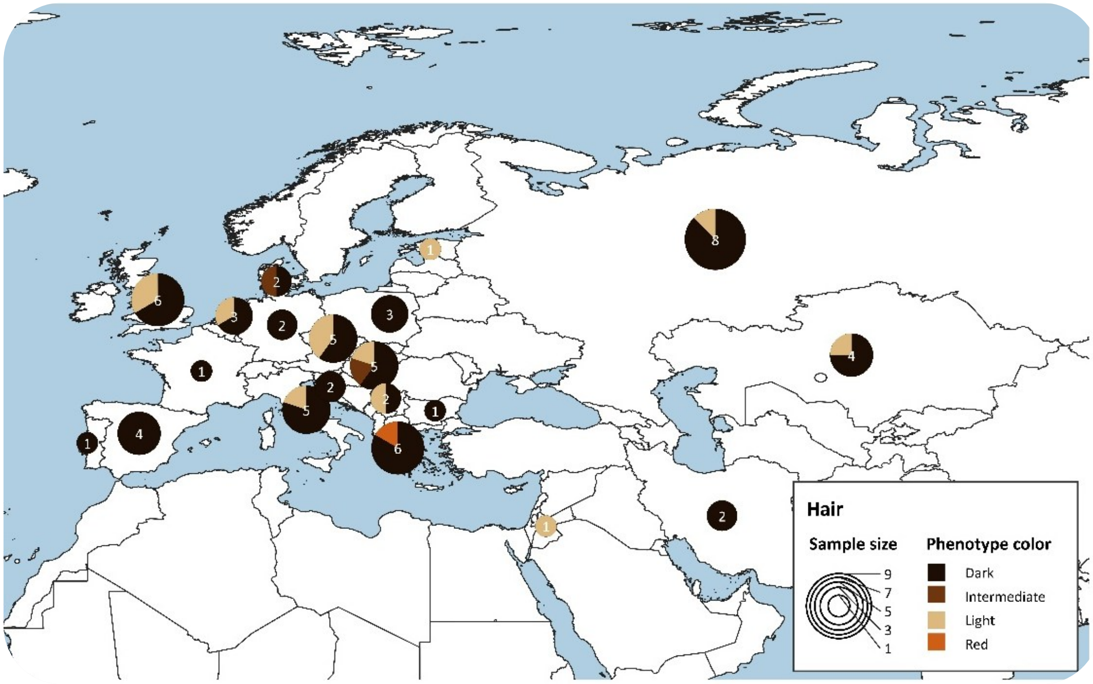


***Temporal and geographical distribution of hair color estimates in Eurasia during the Bronze Age.*** *The map illustrates the spatial distribution of the inferred eye color phenotypes for the Bronze Age period. The size of each pie chart corresponds to the sample size. Eye color results are grouped into 4 categories: Dark (Black, Dark brown/black, and Brown/dark brown estimates), Intermediate (Brown and Dark blond), Light (Blond), and Red.*

### Figure S24

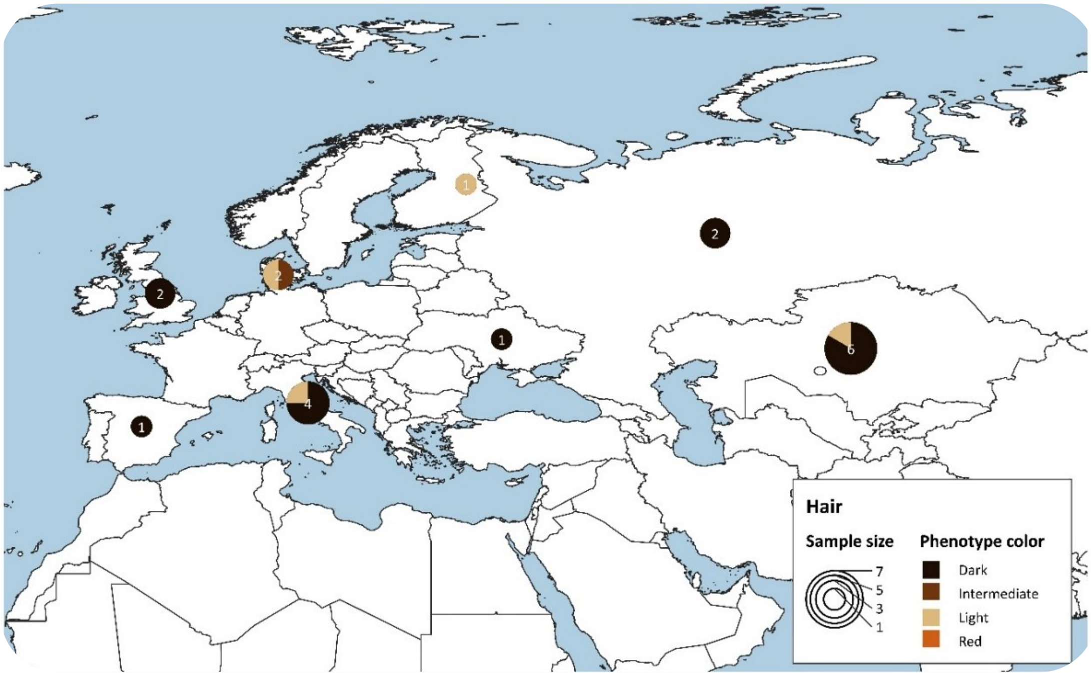


***Temporal and geographical distribution of hair color estimates in Eurasia during the Iron Age.*** *The map illustrates the spatial distribution of the inferred eye color phenotypes for the Iron Age period. The size of each pie chart corresponds to the sample size. Eye color results are grouped into 4 categories: Dark (Black, Dark brown/black, and Brown/dark brown estimates), Intermediate (Brown and Dark blond), Light (Blond), and Red.*

### SI References


1. L. Chaitanya, *et al.*, The HIrisPlex-S system for eye, hair and skin colour prediction from DNA: Introduction and forensic developmental validation. *Forensic Sci Int Genet* **35**, 123–135 (2018).

2. S. Walsh, *et al.*, Developmental validation of the HIrisPlex system: DNA-based eye and hair colour prediction for forensic and anthropological usage. *Forensic Sci Int Genet* **9**, 150–161 (2014).

3. S. Walsh, *et al.*, Global skin colour prediction from DNA. *Hum Genet* **136**, 847–863 (2017).

4. B. Sousa da Mota, *et al.*, Imputation of ancient human genomes. *Nat Commun* **14**, 3660 (2023).

5. R. Nielsen, J. S. Paul, A. Albrechtsen, Y. S. Song, Genotype and SNP calling from next-generation sequencing data. *Nat Rev Genet* **12**, 443–451 (2011).

6. Q. Fu, *et al.*, Genome sequence of a 45,000-year-old modern human from western Siberia. *Nature* **514**, 445–449 (2014).

7. T. Günther, *et al.*, Population genomics of Mesolithic Scandinavia: Investigating early post-glacial migration routes and high-latitude adaptation. *PLoS Biol* **16**, e2003703 (2018).

8. L. Orlando, *et al.*, Ancient DNA analysis. *Nature Reviews Methods Primers* **1**, 14 (2021).

9. Andrews S. (2010), Andrews S. (2010). FastQC: a quality control tool for high throughput sequence data. Available online at: http://www.bioinformatics.babraham.ac.uk/projects/fastqc.

10. M. Schubert, S. Lindgreen, L. Orlando, AdapterRemoval v2: rapid adapter trimming, identification, and read merging. *BMC Res Notes* **9**, 88 (2016).

11. H. Li, R. Durbin, Fast and accurate short read alignment with Burrows–Wheeler transform. *Bioinformatics* **25**, 1754–1760 (2009).

12. D. M. Church, *et al.*, Modernizing Reference Genome Assemblies. *PLoS Biol* **9**, e1001091 (2011).

13. H. Li, *et al.*, The Sequence Alignment/Map format and SAMtools. *Bioinformatics* **25**, 2078–2079 (2009).

14. D. C. Koboldt, *et al.*, VarScan 2: Somatic mutation and copy number alteration discovery in cancer by exome sequencing. *Genome Res* **22**, 568–576 (2012).

15. R Core Team, R: A Language and Environment for Statistical Computing. R Foundation for Statistical Computing, Vienna, Austria. [Preprint] (2021). Available at: https://www.R-project.org/.

16. M. A. DePristo, *et al.*, A framework for variation discovery and genotyping using next-generation DNA sequencing data. *Nat Genet* **43**, 491–498 (2011).

17. M. Byrska-Bishop, *et al.*, High-coverage whole-genome sequencing of the expanded 1000 Genomes Project cohort including 602 trios. *Cell* **185**, 3426-3440.e19 (2022).

18. Picard Tools. Broad Institute. Available at: https://broadinstitute.github.io/picard/.

19. A. S. Hinrichs, The UCSC Genome Browser Database: update 2006. *Nucleic Acids Res* **34**, D590–D598 (2006).

20. H. Li, A statistical framework for SNP calling, mutation discovery, association mapping and population genetical parameter estimation from sequencing data. *Bioinformatics* **27**, 2987–2993 (2011).

21. S. Rubinacci, D. M. Ribeiro, R. J. Hofmeister, O. Delaneau, Efficient phasing and imputation of low-coverage sequencing data using large reference panels. *Nat Genet* **53**, 120–126 (2021).

22. A. McKenna, *et al.*, The Genome Analysis Toolkit: A MapReduce framework for analyzing next-generation DNA sequencing data. *Genome Res* **20**, 1297–1303 (2010).

23. T. M. T. Carratto, *et al.*, Evaluation of the HIrisPlex-S system in a Brazilian population sample. *Forensic Sci Int Genet Suppl Ser* **7**, 794–796 (2019).

24. L. A. Marano, J. D. Andersen, F. T. Goncalves, A. L. O. Garcia, C. Fridman, Evaluation of HIrisplex-S system markers for eye, skin and hair color prediction in an admixed Brazilian population. *Forensic Sci Int Genet Suppl Ser* **7**, 427–428 (2019).

25. D. M. Hohl, *et al.*, Applicability of the IrisPlex system for eye color prediction in an admixed population from Argentina. *Ann Hum Genet* **86**, 297–327 (2022).
